# Engineered Reversible Inhibition of SpyCatcher Reactivity Enables Rapid Generation of Bispecific Antibodies

**DOI:** 10.1101/2023.10.27.564397

**Authors:** Christian Hentrich, Mateusz Putyrski, Hanh Hanuschka, Waldemar Preis, Sarah-Jane Kellmann, Melissa Wich, Manuel Cavada, Sarah Hanselka, Francisco Ylera

## Abstract

The precise regulation of protein function is essential in biological systems, and achieving such control is a fundamental objective in the fields of chemical biology and protein engineering. Here, we describe a straightforward method to engineer functional control into the isopeptide bond-forming SpyTag/SpyCatcher protein ligation system. First, we performed a cysteine scan of SpyCatcher, exchanging each amino acid in the structured region against cysteine. Except for the two known reactive and catalytic residues, none of these mutations abolished reactivity. In a second screening step, we modified the cysteines with disulfide bond-forming small molecules and screened for reactivity again. Here we found 8 positions that, when modified, strongly inhibited reactivity. This inhibition could be reversed by treatment with reducing agents. We call such a reversibly inhibitable SpyCatcher “SpyLock”.

We then used “BiLock”, a fusion of SpyLock and wildtype SpyCatcher, in combination with SpyTagged antibody fragments to generate bispecific antibodies. A first antibody was reacted with the regular SpyCatcher moiety, followed by unlocking of the SpyLock through reduction and its reaction with a second antibody. This method to generate bispecific antibodies requires only a single antibody format and is readily scalable, facilitating the screening of a large number of antibody combinations. We demonstrate the utility of this approach to screen anti-PD-1/anti-PD-L1 bispecific antibodies using a cellular reporter assay.

## Introduction

Biology requires the ability to regulate protein functionality on fast time scales. One common mechanism to do so is allostery, where classically a multimeric protein changes conformation and activity based on the reversible binding of effectors away from the active site^1^. The term allostery has since been extended to include monomeric proteins that are able to change conformation based on effector binding or posttranslational modification. One illustrative example is nitrogen regulatory protein C, a bacterial transcriptional regulator, which changes conformation after phosphorylation^2^.

The engineering of functional control into proteins is an active field of research within basic science and it also has biotechnology applications^3^. One prominent method is the insertion of allosteric domains into other proteins. For example, antibodies allosterically regulated by calmodulin binding proteins have been constructed by inserting calmodulin between the antibody heavy and light chain of an scFv^4^. Another strategy to engineer control is the insertion of a photoswitchable small molecule into the protein, at a position where it perturbs the protein activity through isomerization, thus making protein function controllable by light^5^.

Post-translational modifications are one of the main ways cells regulate protein function^6^. Such posttranslational modifications can naturally occur at cysteines, for example through reversible S-thiolation with glutathione^7^. Glutathionylation of the p50 subunit of NF-κB inhibits binding of this transcription factor to DNA; similarly, glutathionylation of MEKK1 kinase leads to its inactivation^8, 9^.

SpyTag/SpyCatcher is a protein ligation technology in which a small peptide (SpyTag) and a small protein (SpyCatcher) form an isopeptide bond between aspartate 7 in the SpyTag and lysine 31 in the SpyCatcher (Fig. 1A)^10^. The reaction, catalyzed by glutamate 77 in SpyCatcher, is both fast and specific. Two improved versions of SpyCatcher have been engineered, both improving SpyTag-SpyCatcher affinity and accelerating the reaction velocity over the respective previous version^11, 12^. SpyTag/SpyCatcher has proven an extremely versatile technology, with applications for example in vaccine development, nanopore sequencing, or enzyme stabilization^13^. We have previously used this technology to develop a modular antibody platform, in which SpyTagged Fab antibody fragments can be rapidly combined with a wide variety of prefabricated SpyCatcher fusion proteins for site-specific labeling and/or control of oligomeric state and antibody format^14^. We also developed SpyDisplay, a versatile phage display selection technology where SpyTagged antibodies are presented on phages carrying SpyCatcher-pIII fusions^15^.

**Figure 1:**
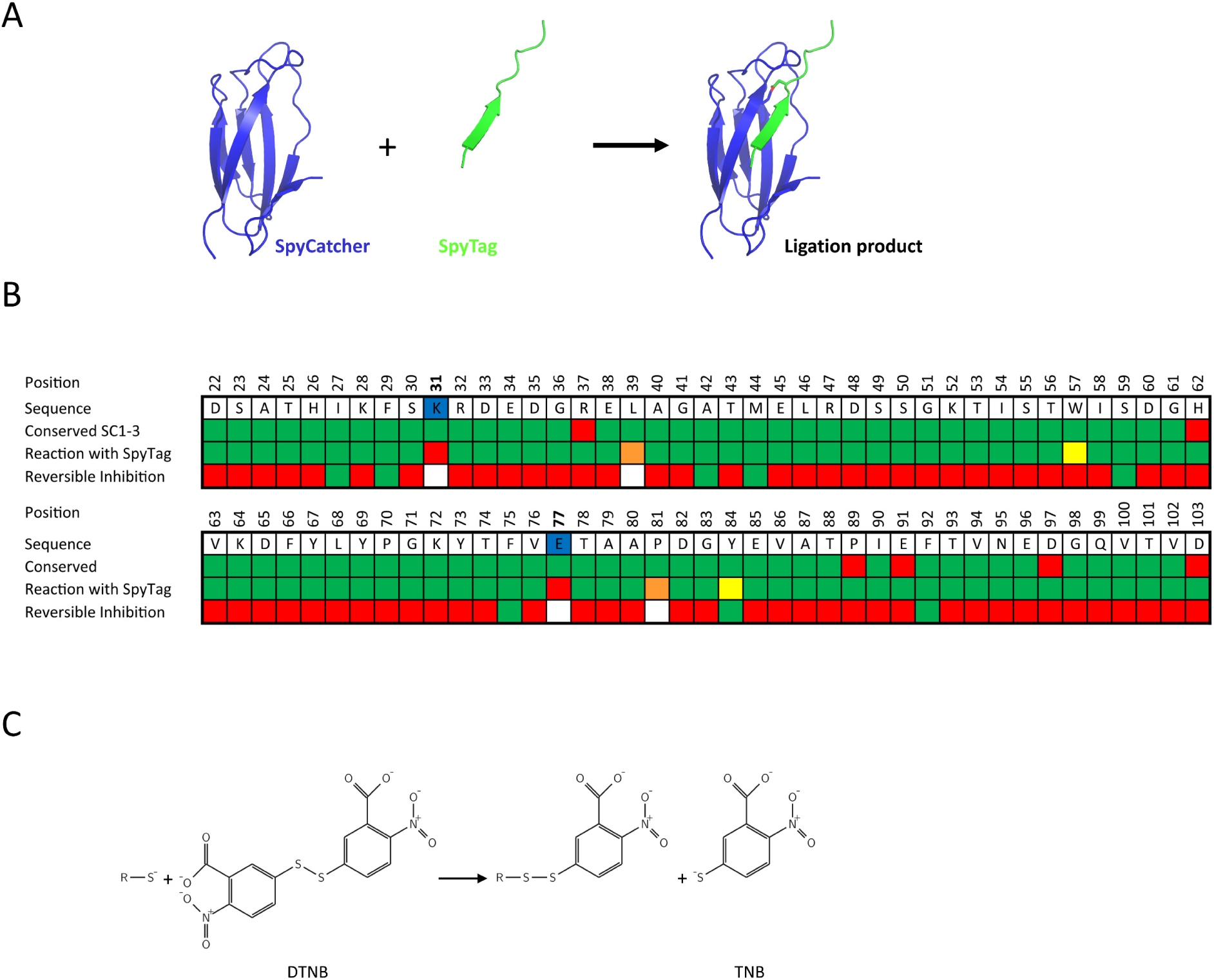
Cysteine scan for reversibly inhibitable SpyCatcher3 mutants. A) Model of the SpyTag-SpyCatcher reaction based on the crystal structure^24^. SpyCatcher (blue) reacts with SpyTag (green), forming a covalent isopeptide bond (red). B) Heat map depicting the cysteine scan of SpyCatcher3 for reversible inhibition upon modification with Ellman’s reagent (DTNB). The first two rows show position and amino acid of SpyCatcher3, with reactive and catalytic residues in blue. The third row signifies amino acid conservation among SpyCatcher1, SpyCatcher2, and SpyCatcher3, using green for conserved and red for differing positions. The fourth row assesses reactivity of unmodified cysteine mutants with SpyTag within 90 minutes: green for fully reactive, yellow for slightly reduced reactivity, orange for strongly reduced reactivity, and red for unreactive. The fifth row evaluates reversible inhibition potential with Ellman’s reagent: green for yes, red for no, and white for not tested. C) Reaction of a deprotonated cysteine residue with Ellman’s reagent (DTNB).

Cysteine scanning mutagenesis is an established method to interrogate protein function. It has been used in the past for example to study the channel forming amino acids of ion pumps and channels, where solvent-exposed thiol groups react much faster with cysteine-reactive probes than buried residues^16^. In other studies, cysteines have been introduced close to the active site of enzymes, in order to conjugate libraries of small molecules for inhibitor screening, where local proximity would also allow the identification of low activity compounds as a starting point for structure-activity relationship studies^17^.

In another study, Yildiz et al. used cysteine scanning to identify an allosteric site in a protease from Dengue virus. 8 amino acids at suspected allosteric sites were mutated to cysteine. One of them, when modified with cysteine-reactive probes, quantitatively inhibited protease activity, and activity was restored after removal of the probe^18^. Harder et al. used a similar approach to block a proton pump with a methanethiosulfonate, which could be removed by reduction^19^.

Our purpose in this study was to engineer functional control into SpyCatcher. Specifically, we wanted to inhibit SpyCatcher in a reversible manner, so that we could activate SpyCatcher reactivity at will. Direct, orthosteric inhibition of SpyCatcher’s active site has been achieved in a straightforward manner by protecting the isopeptide bond-forming lysine 31 with a photocleavable protection group through use of artificial amino acids^20, 21^. We instead aimed to regulate SpyCatcher activity indirectly through S-thiolation of engineered cysteines. SpyCatcher, which is devoid of endogenous cysteines, has no suspected allosteric regulation sites. Therefore, we performed a cysteine scan of every amino acid in its structured region to identify mutants with lowered activity when modified by disulfide-forming small probes. We identified several such positions and found the inhibition to be reversible upon reduction. We call such reversibly inhibitable SpyCatcher mutants “SpyLock”.

We then applied SpyLock for the generation of bispecific antibodies. Bispecific antibodies are typically generated by fusing two distinct antibodies, yielding a single molecule capable of binding different epitopes, often on separate antigens. They have proven to be promising therapeutic agents, particularly in oncology^22^. Production of bispecific antibodies with correct light and heavy chain pairing is not trivial and has led to the development of a plethora of formats for bispecific antibody screening^23^. Utilizing a fusion of SpyLock and regular SpyCatcher, which we call “BiLock”, we rapidly generated bispecific antibodies from SpyTagged Fab fragments and demonstrated their efficacy in cellular assays.

## Results

### Screening for reversibly inhibitable SpyCatchers

In order to engineer functional control into SpyCatcher, we focused on the ordered region of SpyCatcher according to the known crystal structure^24^. We chose SpyCatcher3, the fastest reacting SpyCatcher variant, as being able to control this version should have the highest interest for various applications. We therefore expressed 81 cysteine mutants of every amino acid in SpyCatcher3 between Asp22 and Asp103 in *E. coli*, except for S49C, which is an already described cysteine mutant used for site-specific labeling of SpyCatcher^25^. All cysteine mutants could be expressed and purified, with yields between 3.2 and 17.1 mg/L (median 6.9 mg/L) of purified protein from 50 mL expression cultures.

We then tested the ability of these mutated SpyCatchers to react with MBP (maltose binding protein) fused to SpyTag2. Two residues have been described as essential for the isopeptide formation between the SpyTag and SpyCatcher: Lys31, which is the isopeptide bond-forming amino acid of SpyCatcher, and Glu77, which is a catalytic amino acid for the reaction^10^. As expected, cysteine mutations at these positions completely abolished the reactivity of SpyCatcher3. However, to our surprise, every other SpyCatcher3 mutant remained reactive, albeit in 4 cases (L39C, W57C, P81C, Y84C) with strongly reduced apparent reaction rate (Fig. 1B, Suppl. Fig. 1).

Next, we modified the cysteines of all SpyCatcher3 mutants with Ellman’s reagent (DTNB, 5,51-dithiobis(2-nitrobenzoic acid)), a disulfide bond forming molecule that adds a relatively bulky and charged thionitrobenzoic acid group to the cysteine^26^ (Fig. 1C). We tested reactivity of such modified proteins again and could observe strongly reduced reactivity for 8 SpyCatcher mutants: I27C, F29C, A42C, M44C, S59C, F75C, Y84C, and F92C (Fig. 1B, Suppl. Fig. 2).

Effective control of SpyCatcher function requires this inhibition to be reversible. To test this, we took advantage of the reversible nature of disulfide bond formation and incubated the 8 modified cysteine-SpyCatchers3 with 10 mM TCEP (tris(2-carboxyethyl)phosphine), a common reducing agent. When removing the thionitrobenzoic acid group by reduction and restoring the free cysteine, reactivity with MBP-Spytag2 was restored for all 8 mutants (Fig. 1B, Suppl. Fig. 3).

We thus have identified 8 different mutants of SpyCatcher3 that can be inactivated by reaction with Ellman’s reagent and reactivated by subsequent reaction with a reducing agent and thereby established functional control of SpyCatcher’s enzymatic activity. All 8 identified positions were conserved between SpyCatcher versions 1-3 (Fig. 1B).

In order to test whether this reversible inhibition was particular to Ellman’s reagent or a more general phenomenon, we tested HPDP-biotin (N-[6-(biotinamido)hexyl]-3’-(2’-pyridyldithio)propionamide), which was designed to reversibly label cysteines. Indeed, despite not carrying a charge like TNB (5-thio-2-nitrobenzoic acid), HPDP-biotin was also able to reversibly inhibit the reactivity of the cysteine mutants (Suppl. Fig. 4). The first 5 mutations I27C, F29C, A42C, M44C, S59C were inhibited very strongly by both reagents, indicating that the loss of activity is independent of the charge of TNB but more likely due to the size of the introduced side chain. The last 3 mutations F75C, Y84C, F92C were more strongly inhibited by TNB, thus potentially dependent on charge for inhibition.

### Functional characterization of inhibitable SpyCatchers

Analysis of the 8 abovementioned positions mapped onto the structure of SpyCatcher revealed that none of them protruded towards the SpyTag binding groove (Fig. 2A). Instead, we noticed that 6 out of 8 wildtype residues were rather hydrophobic amino acids, and 5 of these pointed towards the hydrophobic core of the beta-barrel at the center of the SpyCatcher’s Ig fold.

**Figure 2:**
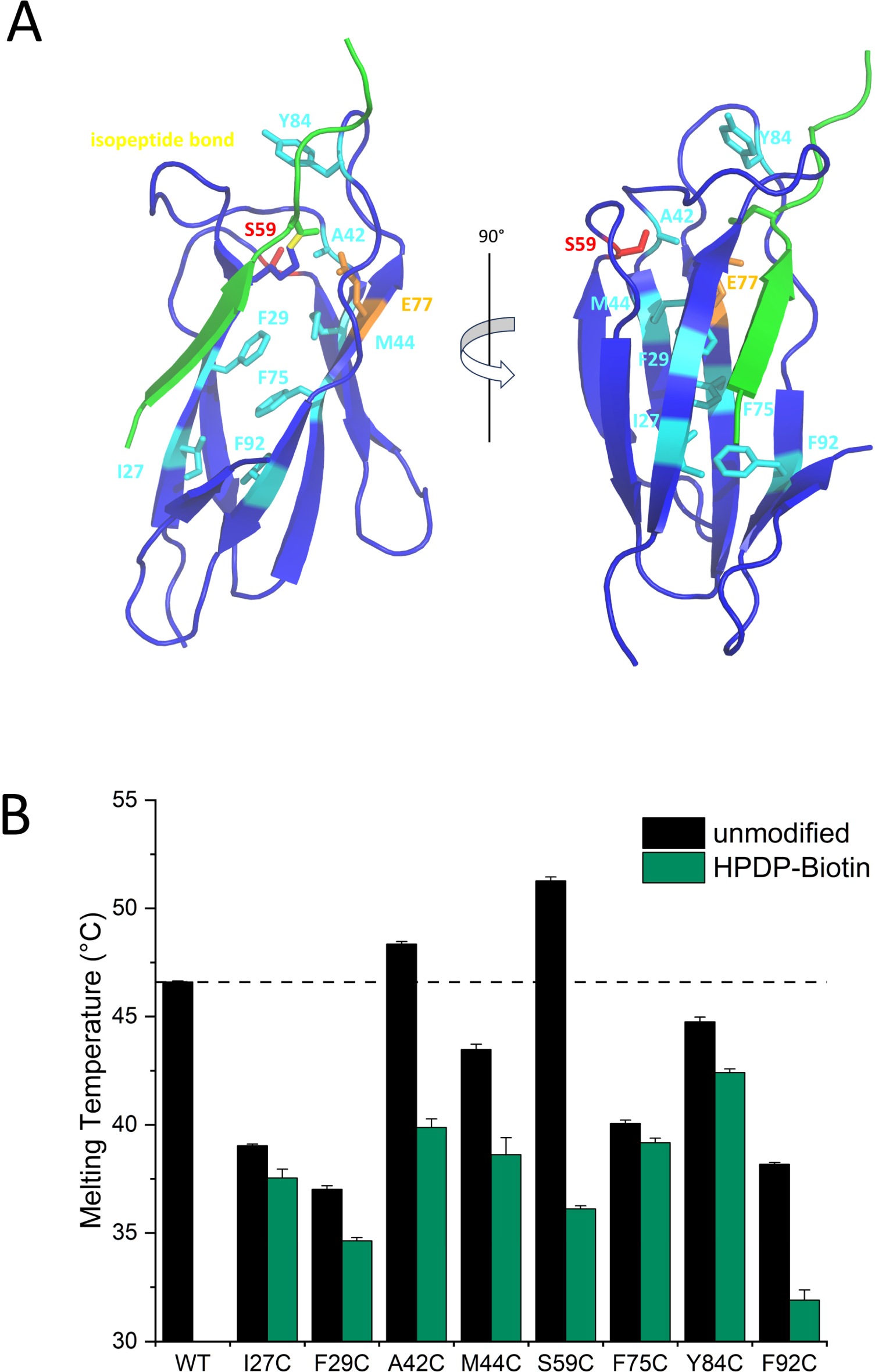
SpyLock-enabling residues within the SpyCatcher structure and the thermal stability of corresponding cysteine mutants. A) Model of SpyCatcher1 (blue) and SpyTag1 (green) with the residues capable of reversible inhibition upon substitution with cysteine highlighted in red (serine 59) or cyan. The catalytic residue is marked in orange, the isopeptide bond in yellow. Two views, rotated by 90 degrees. B) Melting temperatures (determined by nanoDSF) of all identified reversibly inhibitable cysteine mutants in their unmodified and HPDP-biotin-blocked form, in comparison to wildtype SpyCatcher3. Error bars are standard deviations of 3 measurements.

Because of its missing SpyTag beta strand, the melting temperature of uncoupled SpyCatcher is much lower than after coupling with SpyTag^14^. We measured the melting temperature of the 8 identified cysteine mutants in unmodified form and after modification with HPDP-biotin (Fig. 2B). For unmodified SpyCatcher mutants, all variants with cysteine replacing hydrophobic or aromatic amino acids had decreased melting temperatures, as one would expect from modifying amino acids directed towards the hydrophobic core (Fig. 2A). The A42C and S59C mutants were slightly stabilizing. Biotin-modified SpyCatcher mutants were always destabilized compared to the respective unmodified mutant. This was especially pronounced for the two stabilizing mutants A42C and S59C, with the Tm lowered by 8.5°C and 15°C, respectively. However, the fact that a clear Tm was measurable for each mutant clearly demonstrates that both modified and unmodified SpyCatcher mutants remain folded at room temperature. Reaction speeds of all reversible inhibitable mutants were reduced compared to wildtype SpyCatcher3 (Suppl. Fig. 5) but remained sufficient for effective SpyTag coupling at low micromolar concentrations within one hour.

Among the 8 identified reversibly inhibitable SpyCatchers, we selected the S59C mutation for further study. This decision was based on several factors: the S59C mutation is the most conservative, given that serine and cysteine differ by only a single atom; it demonstrated robust reversible inhibition when exposed to Ellman’s reagent or HPDP-biotin (Fig. 3A); the mutation is located outside the protein’s hydrophobic core, minimizing aggregation potential; and it exhibited the highest stability of the mutants as indicated by its melting temperature.

**Figure 3:**
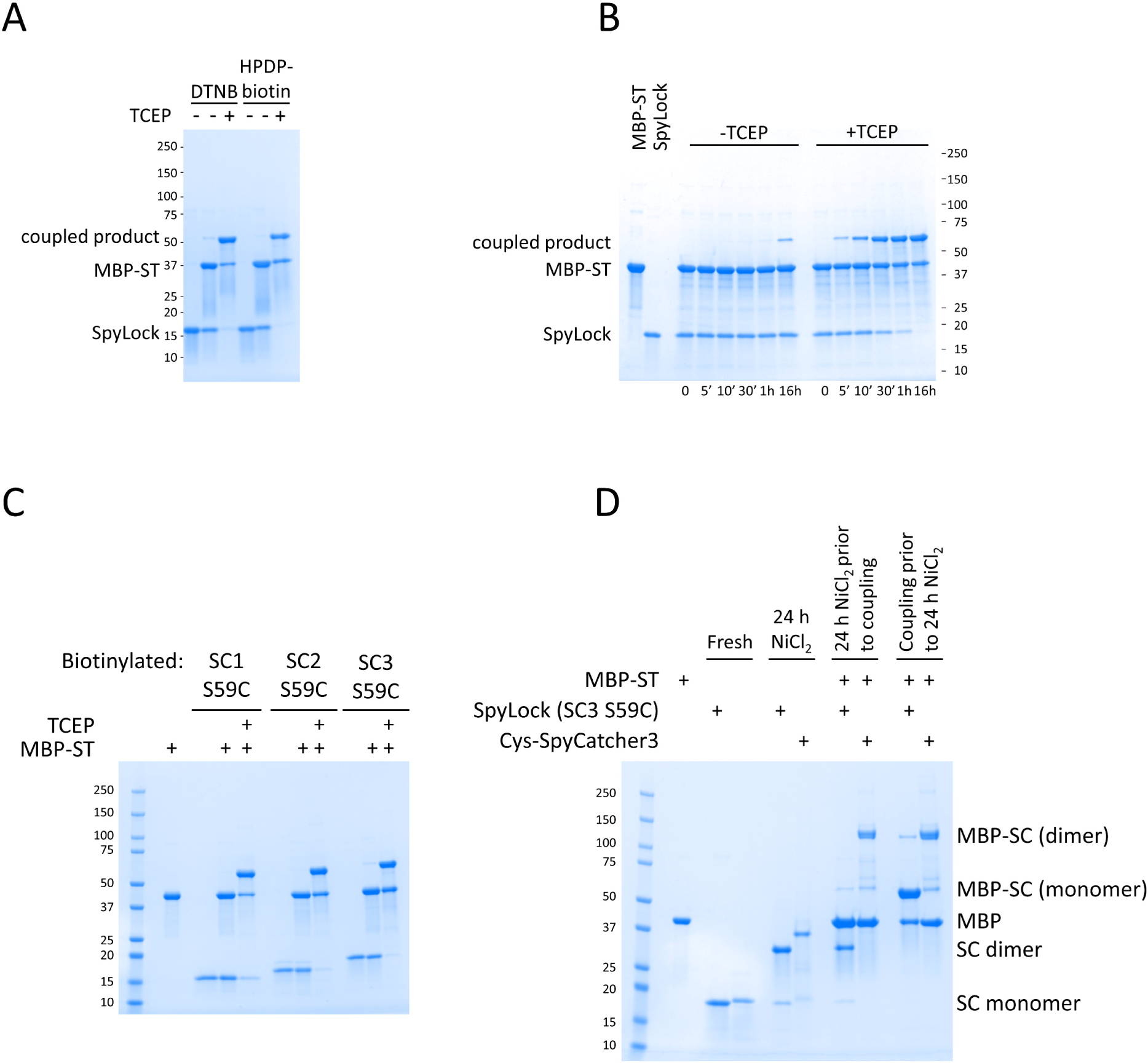
SpyLock coupling reactions. A) Reducing SDS-PAGE gel showing the result of reacting 4 µM SpyCatcher3 S59C (SpyLock) modified with either DTNB or HPDP-biotin was reacted with 6 µM MBP-SpyTag2 in presence or absence of the reducing agent TCEP (5 mM) for 90 minutes. B) Time course of the reaction of 4 µM biotin-SpyCatcher3 S59C (biotin-SpyLock) with 6 µM MBP-SpyTag2 (MBP-ST) in presence or absence of the reducing agent TCEP (5 mM), visualized on a reducing SDS-PAGE gel. C) Reducing SDS-PAGE gel of the S59C variants of SpyCatcher1/2/3 (4 µM) modified with HPDP-biotin, reacting with 6 µM MBP-SpyTag2 for 90 min in presence or absence of the reducing agent TCEP (5 mM). D) Non-reducing SDS-PAGE analysis of NiCl_2_-induced dimerization of SpyLock or N-terminal Cys-SpyCatcher3 and reaction of dimerized catchers with MBP-SpyTag2. Dimerization occurred over 24 h at 37°C. Reaction conditions were identical for pre- and post-dimerization coupling: 10 µM SpyCatcher, 15 µM MBP-SpyTag2, 90 min.

When a reversibly inhibitable SpyCatcher mutant is in its disulfide-modified form, it is in a closed, locked state, preventing function. In contrast, a cysteine mutant of SpyCatcher in its reduced form is akin to an open, unlocked state, allowing it to react freely. Given this functional resemblance to a lock, we have named reversibly inhibitable SpyCatcher mutants ‘SpyLock’. In this manuscript, we usually refer to the S59C mutant of SpyCatcher3 as SpyLock and use the terms “open SpyLock” for the reactive, reduced version with a free thiol group and “closed SpyLock” for the unreactive disulfide version.

We analyzed the time course of coupling reactions of closed SpyLock (treated with HPDP-biotin) in presence and absence of TCEP (Fig. 3B). After one hour, in absence of TCEP, almost no reaction occurred, whereas the reaction was close to completion in presence of the reducing agent. Upon overnight incubation, a small portion of the closed SpyLock reacted also in the absence of TCEP, indicating that rather than fully inhibiting the reaction, the biotinylation of cysteine 59 in SpyLock slows down the reaction by orders of magnitude.

As expected from the fact that all 8 amino acid positions responsible for SpyLock behavior are conserved between SpyCatcher 1, 2, and 3, S59C mutants of SpyCatcher1 and 2 work equivalently (Fig. 3C).

Since proteins with free cysteines tend to dimerize by forming intermolecular disulfide bonds in non-reducing environments, we tested the dimerization potential of SpyLock by incubating open SpyLock with or without MBP-SpyTag2 in PBS/0.5 mM NiCl_2_ for 24 hours. As a control we used SpyCatcher3 with an N-terminal cysteine (Cys-SpyCatcher3). In the absence of MBP-SpyTag2, both SpyLock and Cys-SpyCatcher3 dimerized almost quantitatively. However, there was a strong difference in dimerization status of SpyTag2-coupled Cys-SpyCatcher3 and SpyLock. Cys-SpyCatcher3-SpyTag2 again dimerized almost quantitatively, whereas there was only a very small amount of SpyLock-SpyTag2 dimerization (Fig. 3D). As expected, in contrast to Cys-SpyCatcher3, pre-dimerized SpyLock could not react with SpyTag2 anymore.

Two possible mechanisms could underlie the inhibition observed in S-thiolated, closed SpyLock. The first is strongly reduced SpyTag binding, and the second is an inhibition of the isopeptide bond-forming reaction after binding has occurred. To differentiate between these two mechanisms, we measured the interaction of immobilized SpyTag2 with open and closed SpyLock using biolayer interferometry (BLI) (Fig. 4). During the association phase, SpyCatcher3 and open SpyLock bound to the sensor, while closed SpyLock exhibited only negligible binding, strongly suggesting that it is primarily the SpyTag-SpyCatcher binding that is inhibited. In the dissociation phase of the measurement, no unbinding of SpyCatcher or SpyLock could be detected.

**Figure 4:**
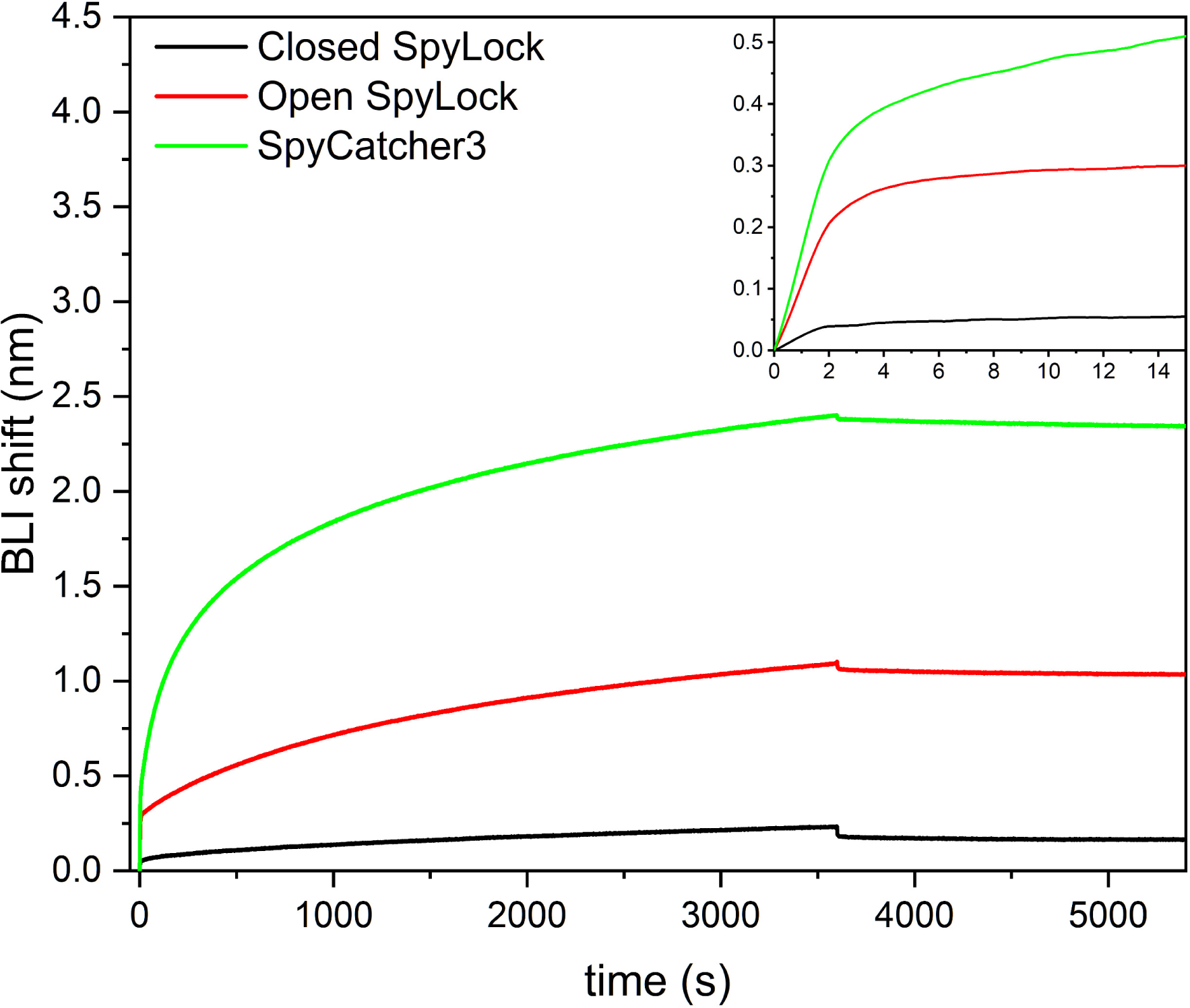
Binding of open and closed SpyLock to SpyTag. BLI sensorgram of SpyLock or SpyCatcher binding to immobilized biotin-SpyTag2. Association until 3600 seconds, followed by dissociation. Inset: Zoom on the first 15 seconds of association.

### SpyLock enables bispecific antibody generation

To demonstrate SpyLock’s usefulness within an application, we chose to employ it for the generation of bispecific antibodies. Therefore, we generated a fusion protein of SpyLock and SpyCatcher, analogous to BiCatcher^14^, which we call “BiLock”. After locking with a disulfide-forming reagent, BiLock has two independent reaction sites for SpyTag, one of which is constitutively active, and one which is inactive and can be unlocked on demand. When a first SpyTagged antibody fragment is added, it will only react with the wildtype SpyCatcher moiety. Then, the SpyLock is unlocked by addition of 5 mM TCEP and a second SpyTagged antibody fragment is added, which can react with the open SpyLock moiety (Fig. 5A, B).

**Figure 5:**
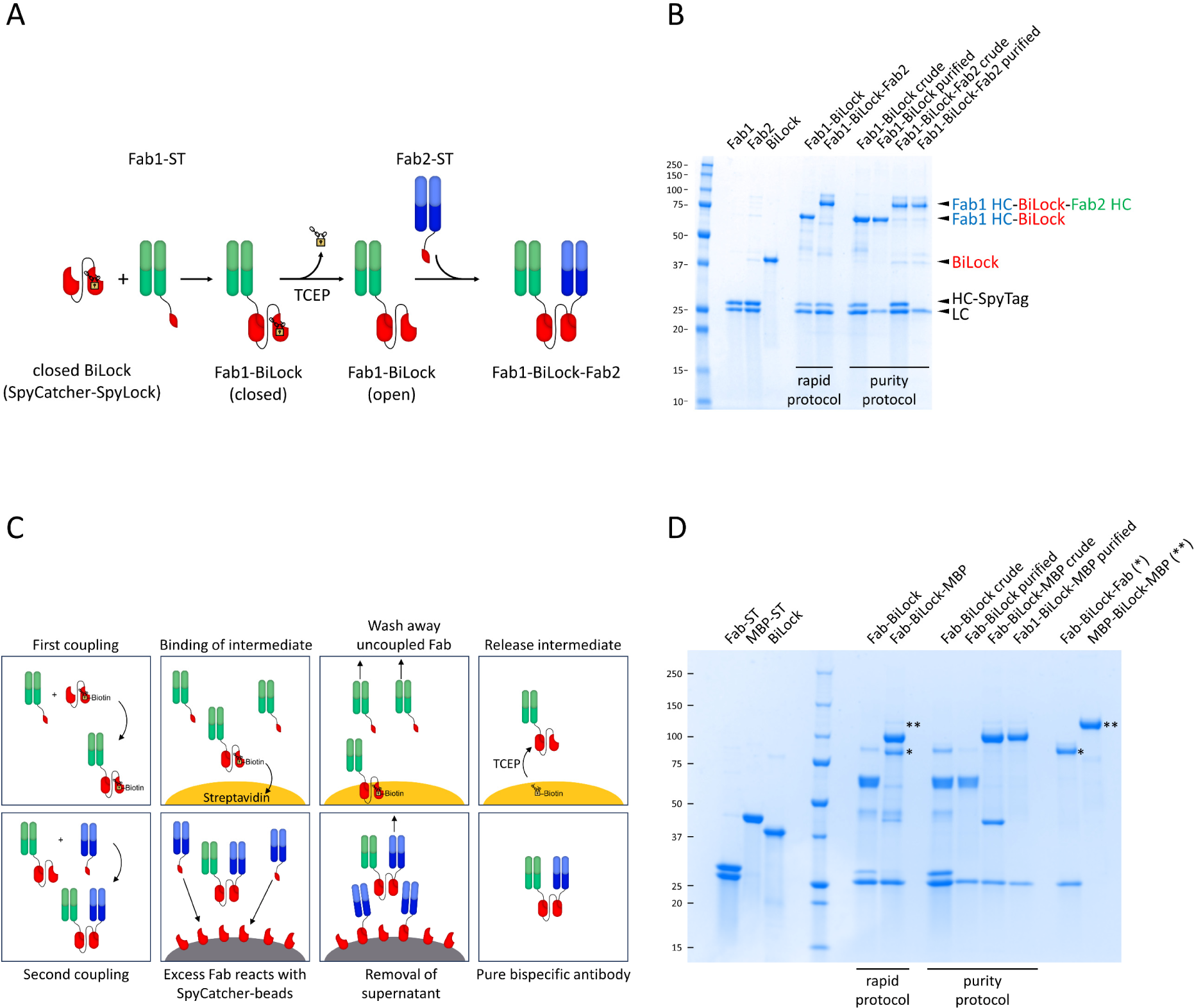
Bispecific antibody generation with BiLock. A) Scheme of the reaction of BiLock (SpyCatcher-SpyLock, red) with two different Fabs (green, blue) to create a bispecific antibody. The disulfide-forming small molecule is signified by a lock. ST is SpyTag and SC is SpyCatcher. B) Reducing SDS-PAGE gel showing sequential generation of bispecific antibodies by either a rapid protocol or a slower protocol involving two purification steps. Rapid protocol: 3 µM BiLock, 3 µM Fab1, 3 µM Fab2, first coupling 45 min. Purity protocol: 3 µM BiLock, 4.5 µM Fab1, 4.5 µM Fab2. First coupling 60 min. Second coupling was carried out overnight in both cases. C) Scheme explaining the purity protocol involving two sequential purification steps. Order left to right, top to bottom. D) SDS-PAGE to assess purity of bispecific antibodies generated with the rapid or purity protocol. Gel analogous to (B), but instead of two Fabs, one SpyTagged Fab and MBP-SpyTag2 were used, allowing to visualize bivalent monospecific contaminants at different apparent molecular weight than the bispecific product. BiLock based on SpyCatcher2 was used to avoid the formation of double bands (Suppl. Fig. 6). Monospecific standards Fab-BiLock-Fab and MBP-BiLock-MBP are loaded in the last two lanes. Corresponding contaminations are indicated by asterisks. Rapid protocol: 10 µM BiLock, 10 µM Fab, 10 µM MBP, first coupling 45 min. Purity protocol: 10 µM BiLock, 15 µM Fab, 15 µM MBP, first coupling 150 min. Second coupling was carried out overnight in both cases.

Using TCEP in the presence of antibodies raises the concern that the intradomain disulfide bonds of the antibodies are also reduced, and the antibodies thereby lose their function. The inter-chain disulfide between heavy and light chains is not required for the Fab stability and is not present in our SpyTagged Fab fragments^14^. To quantify the impact of TCEP, we incubated various Fab fragments with 5 mM TCEP for one week. Utilizing titration ELISA, we observed only minor effects on antigen binding ability (Suppl. Fig. 7). Nevertheless, even though the amount of TCEP present in the samples would therefore not be problematic for most applications, we still decided to routinely remove it by quenching with the non-toxic bis-PEG_3_-azide, as described^27^. This procedure lowers the amount of measurable free TCEP by 97.3%, as determined by use of Ellman’s reagent.

While the abovementioned protocol for assembly of bispecific antibodies is very fast, in practice it is sometimes challenging to achieve very high purity of the bispecific antibodies. This issue arises from the difficulty in determining protein concentrations with sufficient accuracy to ensure precise 1:1 Fab:SpyCatcher coupling. To address scenarios where high purities are essential, we developed a purification protocol that takes advantage of the possibility of using HPDP-biotin with SpyLock (Fig. 5C).

In the first step, we coupled biotin-BiLock with an excess of the first antibody (usually 1.5 molar equivalents) to ensure complete coupling of the SpyCatcher3 within one to three hours. Then, we incubated the product with streptactin or streptavidin beads to pull down the biotinylated BiLock-Fab#1 and washed away excess Fab#1. Any present non-biotinylated BiLock that might have coupled twice to the same antibody is also removed in this step. We then incubated the solid phase with TCEP, thereby simultaneously releasing BiLock-Fab#1 from the solid phase and unlocking the reactivity of the SpyLock moiety. Next, BiLock-Fab#1 was incubated with an excess of Fab#2, also in 1.5-fold molar excess, usually overnight. Afterwards, the bispecific antibodies with excess Fab#2 were incubated with covalently immobilized SpyCatcher3 beads. Unreacted excess Fab#2 reacted with the SpyCatcher3 beads and, after TCEP quenching, the supernatant contained highly pure bispecific antibodies.

The similarity in size between two Fabs makes it impossible to unambiguously demonstrate the purity of bispecific products using gel electrophoresis. To address this, we employed SpyTagged coupling partners of distinct sizes, a Fab and MBP, to create a bispecific MBP-Fab fusion. This allowed for the straightforward differentiation between bivalent monospecific and bispecific products. While there is contamination of the monospecific products visible with the fast protocol, they are essentially absent from the purified samples (Fig. 5D).

### Functional validation of bispecific antibodies generated with BiLock

To validate the functionality of our bispecific antibodies, we performed bridging ELISAs with ocrelizumab and dupilumab as antigens, with one antigen coated on plate and the second HRP-labeled antigen in solution. Our bispecific antibodies, created from two anti-idiotype antibodies, bridge them successfully, while the monospecific bivalent controls do not (Fig. 6A, B).

**Figure 6:**
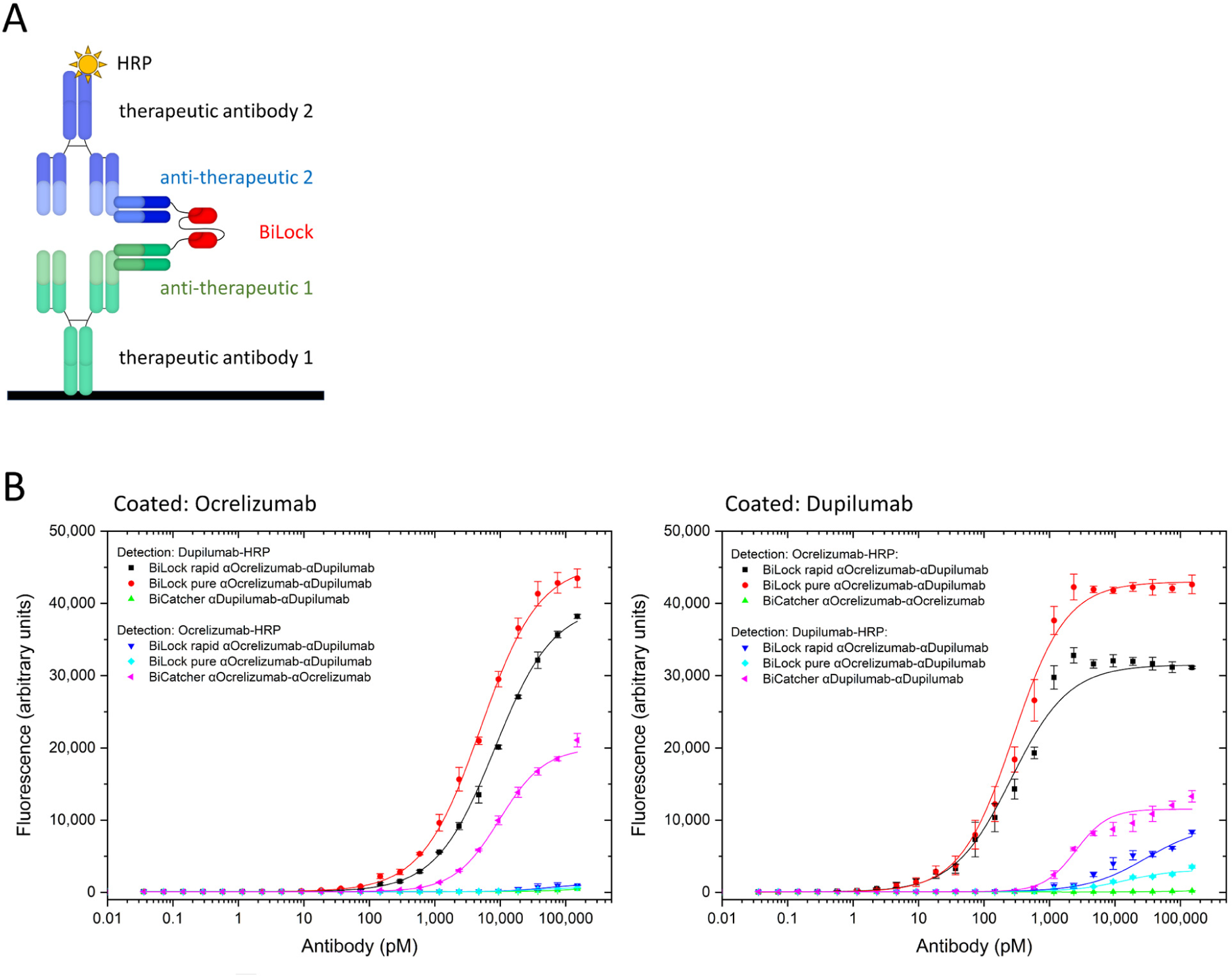
Bispecificity demonstrated by bridging sandwich ELISA. Bispecific anti-ocrelizumab/anti-dupilumab antibodies were produced according to the rapid or purity protocol and their serial dilutions were tested in bridging sandwich ELISA in which antigens dupilumab or ocrelizumab were coated on the plate and HRP-labeled form of either of both antigens was used for detection. A) Schematic representation of the assay. B) ELISA results. Left: ocrelizumab coated. Right: dupilumab coated. Monospecific bivalent controls constructed with BiCatcher3 were also tested with both HRP-labeled antigens. Coating antigen: 1 µg/mL, sandwich titration in 1:2 dilution series starting from 150 nM, HRP-labeled detection antigen: 2 µg/mL. Logistic fit was applied. Error bars represent standard deviation of 3 measurements.

Programmed cell death protein 1 (PD-1) and its ligand PD-L1 are common targets for cancer immunotherapy and interact with a low intrinsic affinity of around 8 µM^28^. Bispecific targeting of PD-1 and PD-L1 has shown enhanced anti-tumor activity in mouse models when compared to combined antibody treatment or individual monotherapies^29^. To ensure compatibility of BiLock-based bispecifics with cellular assays, we generated bispecific antibodies from known therapeutic anti-PD-1 and anti-PD-L1 antibodies and performed a sandwich staining on cells: PD-1-overexpressing HKB11 cells were incubated with bivalent monospecific or bispecific antibodies, then with PD-L1-biotin, and finally were stained with streptavidin-PE followed by flow cytometry. All bispecific constructs stained the cells, whereas none of the bivalent monospecific controls did (Suppl. Fig. 8).

To assess the utility of bispecific SpyLock-based antibodies in cellular screening with more complex assays, we generated 36 bispecific antibodies from 4 anti-PD-1 and 9 anti-PD-L1 antibodies originating from the Pioneer phage display library (Bio-Rad). We used a PD-1/PD-L1 immune checkpoint inhibitor screening assay based on reporter Jurkat-Lucia TCR-hPD-1 cells and Raji-APC-hPD-L1 antigen presenting cells (Invivogen). In this assay, a blockade of the PD-1/PD-L1 interaction results in luciferase expression. All 36 bispecific antibodies exhibited a dose-dependent response, with luciferase signal commencing at a concentration of 10 nM. Notably, some constructs had already reached their maximal signal level at this concentration. (Fig. 7A).

**Figure 7:**
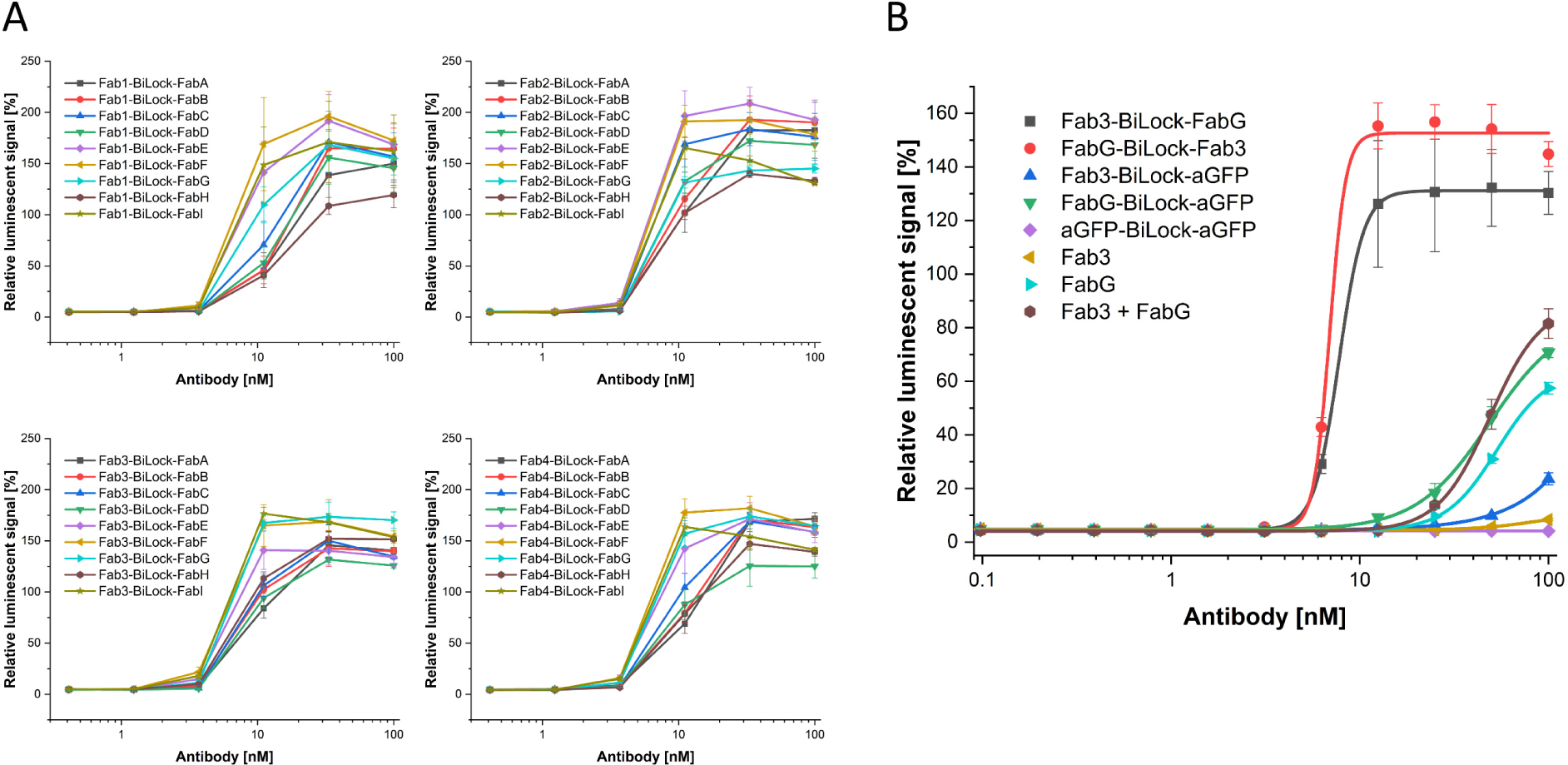
Bispecific antibodies in cellular bioluminescent PD-1/PD-L1 blockade assay. A) 4 anti-PD-1 antibodies (Fab1-4) and 9 anti-PD-L1 antibodies (FabA-I) derived from the Pioneer library were used to construct 36 bispecific antibodies (purity protocol) and tested in PD-1/PD-L1 blockade assay. Results are shown in separate graphs corresponding to 4 anti-PD-1 antibodies tested. B) PD-1/PD-L1 blockade assay performance of one bispecific antibody in comparison to individual Fabs, an equimolar mixture of Fabs (each at the indicated concentration), and additional controls with an irrelevant antibody (anti-GFP). For the anti-PD-1/anti-PD-L1 bispecific antibody, both coupling orders were tested. Logistic fit was applied. All measurements (in A and B) were conducted in triplicate and are presented as the mean ± SD, expressed as a percentage of the signal generated by treating the cells with 50 nM dostarlimab.

We characterized the assay performance of one anti-PD-1/anti-PD-L1 bispecific pair in greater detail (Fig. 7B). We created the bispecific antibody in both possible reaction orders, i.e., coupling either the anti-PD-1 or the anti-PD-L1 antibody first. As controls, we included a monospecific bivalent anti-GFP antibody created with SpyLock-SpyCatcher fusion, anti-GFP/anti-PD-1 or anti-GFP/anti-PD-L1 bispecific constructs, as well as monovalent Fabs. In monovalent Fab format, the anti-PD-1 antibody led to a very low response, while the anti-PD-L1 antibody elicited a modest response at high concentrations. However, in bispecific format, the assay response was very robust, with signal plateau already reached at 10 nM as in the initial screening experiment. The mixture of the two Fabs gave a roughly additive response compared to the individual Fabs. In summary, the data show a strong inhibitory effect of the bispecific antibodies compared to their corresponding Fabs.

The bivalent BiLock-based anti-GFP monospecific control did not yield a detectable signal. However, the bispecific anti-PD-1/anti-GFP or anti-PD-L1/anti-GFP gave slightly higher responses than their monovalent equivalents. This could be due to the fact that the larger constructs are better able to sterically block the PD-1/PD-L1 interaction on the assay cells.

For practical applications, the ability to produce and store large batches of closed BiLock over extended periods would be advantageous. To explore this possibility, we conducted accelerated stability studies on BiLock closed with either Ellman’s reagent or HPDP-biotin. The BiLock modified with Ellman’s reagent demonstrated instability, reverting from its closed state within just two weeks of incubation at 20°C. In contrast, modification with HPDP-biotin maintained a stable closed state for over 16 weeks at the same temperature (Suppl. Fig. 9).

## Discussion

We have employed an easy and straightforward strategy to engineer functional control into SpyCatcher. It consists of conducting a cysteine scan, i.e., producing a cysteine mutant for each amino acid, and testing for functionality before and after modification with a disulfide-forming reagent. We expect this approach to be adaptable to other split proteins like SnoopCatcher, DogCatcher, or split GFP 1-10/11^30, 31, 32^. These proteins are likely very suitable for the engineering-in of reversible inhibition by cysteine scanning, since usually inaccessible amino acids are exposed to solvent in split proteins and can readily react with disulfide-forming reagents.

Besides the known catalytic residue and the isopeptide bond-forming residue, we did not find a single cysteine mutation that abolished enzymatic function of SpyCatcher. However, we did uncover 8 positions that did severely inhibit activity upon modification with a disulfide bond-forming small molecule.

Upon examining these inhibiting positions, it becomes evident that the majority of them are mutations of hydrophobic amino acids that form the hydrophobic core of the Greek-key beta barrel of SpyCatcher’s immunoglobulin fold. This shows that a cysteine oriented towards the hydrophobic core retains the ability to react with small molecules, probably facilitated by the fact that one beta strand of the barrel, the SpyTag, is missing. Introducing bulky or charged modifications at these positions in all probability disturbs the relative orientation of the beta strands and thereby disturbs the reaction center. Instead of leading to aggregation, this disturbance at the hydrophobic core is fully reversible and reduction of the modified cysteine fully restores reactivity of SpyCatcher. None of the amino acids at positions that are capable for generating SpyLocks have side chains directly pointing towards the binding groove. This analysis supports the notion that the engineered regulation is allosteric, affecting the overall structure of SpyCatcher rather than directly inhibiting the active site. Indeed, the serine 59 is one of the inward directed amino acids located furthest away from the groove (Suppl. Fig. 10), yet still strongly abolishes binding of SpyTag when the corresponding cysteine mutant is modified.

Reasoning that more conservative mutations should be beneficial with regards to long-term stability, we decided to focus on the serine 59 to cysteine SpyLock, located in a loop above the beta barrel, at the same side as the reaction center. Already in a reduced state, with just one atom of difference, the reaction speed of this variant is slower compared to that of the wild-type SpyCatcher. This suggests that the geometry of the loops above the beta barrel of SpyCatcher is crucial for reactivity. In the structure of SpyCatcher, serine 59 is oriented inward, and this positioning is very likely preserved in its cysteine mutant form, SpyLock. This inward orientation conceals the sulfhydryl group after coupling with SpyTag, accounting for the slow rate of dimer formation compared to proteins with more exposed cysteine residues. This property is highly desirable in SpyLock, preventing uncontrolled dimer formation and thereby maintaining the intended state of the reagent. Further detailed structural studies would be needed for a comprehensive understanding of the inhibition mechanism. However, for SpyLock with the S59C mutation, evidence from both BLI and nanoDSF data converge to suggest a mechanism of action. Specifically, BLI data show abolished SpyTag binding after TNB modification, while nanoDSF reveals a significantly reduced melting temperature following HPDP-biotin modification. These findings collectively imply that modifying the cysteine with a small molecule disrupts the SpyCatcher structure, maintaining it in a partially unfolded state and thereby abolishing SpyTag binding.

Efforts to control SpyCatcher reactivity have been developed in the past. One straightforward alternative way to inhibit SpyCatcher reactivity is to modify the reactive lysine of SpyCatcher. This has been demonstrated particularly with photocleavable protection groups^20, 21^. While this allows very elegant applications, the SpyLock approach is much easier to implement and more economical to scale as it does not require an expression system capable of implementing artificial amino acids. It can be produced in bacteria and reacted with inexpensive small molecules in bulk. Light-controlled reactivity in general is extremely powerful in specific research settings, but it is not well adaptable to most routine laboratory workflows.

Other strategies using disulfides and redox potential for controlling the reactivity of SpyCatcher have been published, however, these approaches differ fundamentally from our work^33, 34^. Matsunaga et al. established a system based on split-Spy0128, which can be considered a predecessor of the SpyTag-SpyCatcher system^34^. Their approach relies on holding a non-reactive SpyTag-like peptide in the SpyCatcher-like binding pocket with a disulfide bond, thus shielding the access to the reaction center. Upon reduction of the disulfide, the shielding peptide can be displaced by wildtype SpyTag-equivalent peptide and the reaction can occur. Wu et al. have developed a way to change the conformation of SpyTag in such a way that it is strained and cannot react anymore^33^. Upon cleavage of a disulfide bond, the SpyTag can relax and react with SpyCatcher. In yet another example, SpyTag incorporated into a light-sensitive protein remains unreactive in the dark and becomes reactive after unfolding of the protein through irradiation with blue light^35^. All these approaches are highly engineered specialized applications, whereas our SpyLock approach relies only on a single engineered cysteine and is probably generalizable to other split proteins.

Another interesting approach based on a non-reactive SpyTag has recently been developed by the Howarth laboratory^36^. Here, instead of using a disulfide, a non-reactive SpyTag is held in place by a protease-cleavable linker. Upon cleavage by a protease, the non-reactive SpyTag dissociates and unmasks SpyCatcher reactivity.

A notable limitation of SpyLock is its reliance on purified proteins, rendering it incompatible with *in vivo* use. While it may be feasible to expand the concept to SpyCatcher constructs containing two cysteines that in the closed state form an intramolecular disulfide bond, such an approach would require extensive engineering and the activation of the molecule by the disulfide bond reduction would be dependent on local redox potential. In this study, we have purified each cysteine mutant for the cysteine scan. It may however also be possible to perform an *in vitro* selection combined with next generation sequencing to identify reversibly inhibitable split proteins, similar to techniques used for binding interface mapping^37^.

In this work, we used Ellman’s reagent and HPDP-biotin to generate the disulfide required for inhibiting the reactivity of cysteine mutants of SpyCatcher. Depending on the position of the cysteine, we could see differences in locking efficiency. More noticeable was the stability of the disulfide bond over time. Whereas BiLock blocked with HPDP-biotin remained stable, BiLock-TNB was unstable over time and therefore not suitable for bulk preparations. We expect that many disulfide-forming small molecules and also peptides would be suitable to inhibit SpyLock reactivity. Besides biotin, activated variants of other small molecules compatible with their cognate affinity purification reagents such as maltose, azide, alkines, or glutathione can be envisioned. Also activated thiol-reactive peptide tags (e.g. 3-nitro-2-pyridinesulfenyl-activated Strep-tag) could be an option. However, broad availability and low cost are good arguments for the molecules we have chosen in this study.

A straightforward application of SpyLock is the generation of bispecific antibodies. The therapeutic potential and efficacy of bispecific antibodies is widely acknowledged^22^. What all bispecific antibodies have in common is that the screening space grows quadratically with the number of antibody sequences to be tested in combination. As most final therapeutic bispecific antibody formats are labor intensive to produce, it is useful to reduce the number of bispecific antibodies that need to be converted into the final format. Therefore, a first screening step in an alternative format for general functionality can massively reduce the workload and accelerate candidate selection. To this end, many different approaches have been developed, using protein ligation technologies such as split inteins, sortase A, transglutaminase, bispecific Fab-SpyCatcher-Fc fusion proteins, or SpyCatcher-SnoopCatcher fusions^23^.

Our SpyLock strategy has several key differences from most of these approaches. The biggest advantage of using a SpyLock-SpyCatcher dimer is the ability to use only a single antibody format – there is no asymmetry in the antibody fragments used, only SpyTagged antibodies are required, and they can be produced rapidly from bacteria^14^. Each antibody can be coupled as the first or as the last, and each antibody is compatible with any further antibody potentially to be screened in the future. Screening for biparatopic antibodies from a single selection campaign is also easily possible. This advantage is especially pronounced when the antibodies are directly generated with a SpyTag, such as we have established for Pioneer and HuCAL antibody libraries^15, 38^, but it is easy to reclone any human antibody fragment for SpyTagged expression in *E. coli*. Alternatively, SpyTagged antibody fragments can be expressed in eukaryotic cell culture. Another strength of our approach is the possibility to perform the bispecific antibody generation in two ways. Our rapid approach only involves the sequential mixing of few reagents and can be performed within 90 minutes. Its purity in practice is limited by the accuracy of antibody concentration determination. Too much of the first antibody leads to a byproduct of antibody #1 dimers, too little to a byproduct of antibody #2 dimers. Too little of the second antibody leads to monomeric antibody #1 as a byproduct, too much to monomeric antibody #2 as impurity. In the context of very high throughput applications, these might be acceptable compromises.

Alternatively, due to the presence of biotin on the closed SpyLock, we are able to couple with excess of antibody #1, drive the reaction to completion and purify away any unreacted antibody #1 monomers as well as antibody #1 dimers. The second coupling can also be carried out in excess of antibody #2, and leftover antibody #2 can easily be removed due to its reactive SpyTag. This procedure strongly increases the purity of the bispecific antibody, and, while more complex, can also be scaled up and automated easily. This purification protocol allows a more efficient use of monomeric Fabs compared to Driscoll et al., which requires a 5-fold Fab excess in the second coupling step^36^. The amount of Fab required becomes an important economical factor when producing a large number of bispecific antibody pairs. However, the introduction of a biotinylated cysteine or Avi tag^39^ in the protease sensitive linker applied by Driscoll et al. would allow the use of our two-step purification protocol.

SpyLock for generation of bispecific antibodies is open to implementation in many different geometries. As demonstrated by Driscoll et al., the precise geometry of the catcher-based bispecific antibodies can have a profound impact on the functionality of the reagents^36^. This again exemplifies the necessity of final verification of the selected antibody pair in the format in which the bispecific antibody will be therapeutically administered. We chose here a construct analogous to BiCatcher^14^, with a flexible linker sequence. This linker is readily adjustable in terms of length and stiffness, enabling the exploration of various geometries and mimicking the typical paratope reach distance of different antibody formats. To further expand the set of SpyLock-based bispecific formats for screening, it should also be possible to combine SpyLock with knob-in-hole Fc^40^ domains, with the caveat that exposed disulfides in the hinge region should either be avoided or, alternatively, reoxidized after bispecific generation. Less canonical formats, such as SpyCatcher-Fc-SpyLock for tetravalent bispecific antibodies should also be easy to implement. SpyLock significantly expands the toolbox for the modular construction of antibodies based on SpyTag/SpyCatcher, providing a straightforward pathway to bispecific antibodies. We expect that the introduction of redox control of protein ligation through engineered reversible inhibition will prove useful in other fields.

## Methods

### Construct design and cloning

DNA sequences of the cysteine mutants of SpyCatcher and the SpyLock-SpyCatcher fusion constructs were ordered from Twist Bioscience either as gene strands, followed by Gibson assembly into pET-28a(+), or as genes cloned into pET-28a(+). SpyCatcher mutant positions are shown in Figure 1B.

SpyCatcher sequences have been described previously^10, 11, 12^. The BiLock sequence is identical to previously described BiCatcher sequence, with an S59C mutation in the second SpyCatcher^14^. For the coupling purity assay, BiLock based on SpyCatcher2 was used to avoid the appearance of double bands that occur with SpyCatcher3 (Suppl. Fig. 6). Plasmids were transformed into BL21 cells for expression.

### Fab and SpyCatcher expression

Expression of Fabs and SpyCatchers was performed as described^14^.

### SpyCatcher3 beads

SpyCatcher3 was immobilized on Profinity Epoxide Resin (Bio-Rad) according to the manufacturer’s instructions. In short: 1.9 g of dry resin was weighed out and allowed to swell in an excess of coupling buffer (50 mM boric acid, 0.5 M K_2_SO_4_, pH 9.0 with NaOH). 97 mg of SpyCatcher3 at 6 mg/mL in coupling buffer was added to 12 mL of settled resin and mixed for 22 hours at room temperature. The coupled resin was washed with PBS and remaining active groups were blocked by addition of 40 mL of 1 M ethanolamine, pH 9.0 and 5 hours incubation at room temperature. After washing with PBS and 100 mM sodium phosphate, 500 mM NaCl, pH 6.0, and subsequently with 100 mM sodium phosphate, 500 mM NaCl, pH 8.0, the resin with 7 mg/mL immobilized SpyCatcher3 was stored in PBS at 4°C.

### Screening for reversible inhibition

In the initial screening phase, we incubated SpyCatcher cysteine mutants at a concentration of 4 µM and MBP-SpyTag2 at a concentration of 6 µM in PBS at room temperature for one hour to evaluate their reactivity. Samples were then boiled in SDS sample buffer, followed by SDS-PAGE. In the second screening step, SpyCatchers were incubated with 2.5 mM TCEP for 1 hour to fully reduce the cysteines. This was followed by buffer exchange with PD MiniTrap G-25 columns (Cytiva) into 100 mM sodium phosphate buffer, pH 8.0. Resulting protein concentrations were determined by A280 measurements. Subsequently, all SpyCatchers were incubated for 1.5 hours at room temperature with 430 µM DTNB (Thermo, from a 10 mM stock in 100 mM sodium phosphate, pH 8.0), equivalent to a tenfold molar excess of the highest concentrated SpyCatcher sample. To assess reactivity of the modified SpyCatchers, 4 µM of TNB-reacted SpyCatchers were incubated with 6 µM MBP-SpyTag2 for 1.5 hours at room temperature, followed by SDS-PAGE. In the third screening step, mutant SpyCatchers that showed strong inhibition when modified with TNB were tested for reversibility of the inhibition. 4 µM TNB-SpyCatchers were incubated for 1.5 hours with 6 µM MBP-SpyTag2 in presence or absence of 10 mM TCEP (sufficient to reduce any excess DTNB) and analyzed by SDS-PAGE.

### Generation of closed SpyLocks

SpyLock constructs (including BiLock fusions) were incubated in PBS/2.5 mM TCEP for one hour, followed by buffer-exchange (MiniTrap G-25 columns, Cytiva) into DTNB-labeling buffer (100 mM sodium phosphate, 1 mM EDTA, pH 8.0) or biotin-labeling buffer (PBS pH 7.4, 1 mM EDTA). Fresh stock solutions of labeling reagents were prepared, 10 mM DTNB in 100 mM sodium phosphate, pH 8.0 or 4 mM EZ-Link HPDP-biotin (Thermo) in DMSO. A tenfold excess of labeling reagent was added to the SpyLock constructs and the labeling reaction was typically carried out overnight at room temperature. The next day, the buffer was exchanged into PBS and the SpyLock concentration was measured using a Bradford assay (Bio-Rad), using BSA as calibration standard. Modified SpyLocks were aliquoted and stored at −80°C for long-term storage, or −20°C for short-term storage.

### TCEP quantification

A standard curve was generated using known TCEP concentrations. The standard and samples were reacted with Ellman’s reagent (Thermo) and photometrically measured, both in accordance with the manufacturer’s instructions. TCEP concentrations in the samples were then calculated using linear regression based on this standard curve.

### SpyLock-SpyTag BLI measurements

For determination of SpyLock-SpyTag interaction, BLI measurements were performed on an Octet RED384 instrument (Sartorius). SpyTag2 peptide with an N-terminal biotin (Intavis) was immobilized on streptavidin sensors (SA, Sartorius) in PBS with immobilization levels of 2 ± 0.5 nm. Quenching was carried out with 10 µg/mL biocytin in PBS. After setting the baseline in PBS/0.1% BSA/0.02% Tween 20/500 mM NaCl, association of SpyLock, SpyLock-TNB, SpyCatcher3 (100 µg/mL in PBS/0.1% BSA/0.02% Tween 20/500 mM NaCl) was assessed for 1 hour, followed by dissociation in PBS/0.1% BSA/0.02% Tween 20/500 mM NaCl for 30 min For reference, sensors were coated with biocytin only. Data were analyzed with Octet Analysis Studio software 12.2.

### Rapid bispecific antibody assembly

For the rapid bispecific antibody assembly an equimolar amount of closed BiLock and the first Fab-SpyTag2, both in PBS, were incubated for 30 minutes at room temperature in PBS. Then, 5 mM TCEP and an equimolar (regarding BiLock) amount of the second Fab-SpyTag2 were added and allowed to react for at least one hour at room temperature. This was followed by addition of 50 mM bis-PEG_3_-azide (Lumiprobe, from undiluted stock) for TCEP quenching if so desired.

### Bispecific antibody assembly with purification

In this approach, the initial coupling of closed biotin-BiLock and the first Fab-SpyTag was carried out with 1.5-fold molar Fab excess for 1-3 hours. Constructs were then added to pre-washed (PBST) and pre-blocked (5% BSA in PBST) magnetic streptavidin beads (Dynabeads M-280, Thermo) or Streptactin-agarose (IBA Lifesciences), with the amount of beads used being tenfold greater than what would be theoretically required based on the manufacturer’s listed binding capacity. Beads were incubated for 1 hour on a rotator, washed 3 times with PBST and incubated with 5 mM TCEP in PBS to release the SpyLock from the beads and at the same time open it. For larger batches of bispecific antibodies, since the first coupling is not yet in quadratic space, this is feasible even for large numbers of antibodies. With calculated concentrations based on the observation that the biotin pulldown and disulfide reduction are quantitative, the supernatant was split into equal parts for the second Fab coupling. The second antibody was added in 1.5x molar excess of the intermediate construct and coupled for at least 3 hours, typically overnight. This was followed by addition of 50 mM bis-PEG_3_-azide for TCEP quenching. To remove excess Fab 2, the bispecific antibodies were incubated with pre-washed SpyCatcher3 beads for 15 minutes and the purified bispecific antibody was collected in the flow-through.

### Sandwich ELISA

Capture antigens (ocrelizumab, dupilumab) were coated on 384 well plates (Nunc Maxisorp MTP, Thermo) at a concentration of 1 µg/mL in PBS, 20 µL/well. The next day, the plates were washed 3x with PBST (PBS with 0.05% Tween 20 (Merck Millipore)) and then blocked with 100 µL of 5% BSA (Sigma-Aldrich) in PBST for 1.5 hours at room temperature and washed again 3x with PBST. Next, 20 µL of a 1:2 dilution series of the bispecific antibodies, starting at a concentration of 15 µg/mL, were transferred onto the plate and allowed to incubate for 1 hour at room temperature. After incubation, the plates were washed five times with PBST before the addition of HRP-conjugated detection antibodies (ocrelizumab-HRP and dupilumab-HRP) with 20 µL/well and concentration of 2 µg/mL, prepared in HISPEC assay diluent (Bio-Rad). HRP conjugates were prepared using the Lynx HRP conjugation kit (Bio-Rad) according to manufacturer’s protocol. Plates were incubated for an additional hour at room temperature. Subsequently, the plates were washed ten times with PBST, preceding detection with QuantaBlu fluorescence detection reagent (Thermo).

### PD-1/PD-L1 flow cytometry assay

HKB11 PD-1 cells were used as stable pool generated by transfecting HKB11 cells with pMAX vector^41^ encoding human PD-1 (UniProt Q15116) and selection with 0.8 mg/mL geneticin (Invivogen).

HKB11 cells (wildtype and PD-1 stable pool) were cultivated in MAC1.0 v110217 medium (Gibco) with 10% heat inactivated FBS (Gibco) and, for the PD-1 overexpressing cells, 0.8 mg/mL geneticin (Invivogen) in 125 mL Optimum Growth flasks (Thomson), shaking at 120 rpm at 37°C in humidified atmosphere containing 5% CO_2_.

For the flow cytometry assay, 3×10^4^ cells/well/40 µL were incubated with 5 nM of mono- or bispecific antibody constructs in flow buffer (DPBS w/o Ca and Mg (PAN-Biotech) with 3% heat inactivated FBS (Gibco)) for 1 hour at 4°C in a V-Bottom 384 well plate (Greiner). After washing twice with 40 µL/well flow buffer, the cells were incubated in flow buffer with 1 nM biotinylated human PD-L1 (ACRO Biosystems) for 30 min at 4°C. Following incubation, cells were washed twice with 40 µL/well flow buffer and stained with streptavidin-PE (Qiagen) for 30 min at 4°C. Cells were washed twice in 40 µL/well flow buffer and resuspended in 40 µL/well flow buffer with 2.5 µg/mL DAPI (Merck). The median fluorescence intensity of living single cells was determined in triplicates using a ZE5 Cell Analyzer (Bio-Rad) and FCS Express (De Novo Software) for analysis.

### PD-1/PD-L1 immune checkpoint cellular assay

Jurkat-Lucia TCR-hPD-1 effector cells and Raji-APC-hPD-L1 antigen presenting cells were obtained from Invivogen as components of the PD-1/PD-L1 Bio-IC assay (Invivogen, #rajkt-hpd1). The cells were cultivated in IMDM with glutamine and HEPES (Gibco) with 10% heat inactivated FBS (Gibco) in T75 or T175 flasks (Sarstedt) at 37°C in humidified atmosphere containing 5% CO_2_. Growth medium was supplemented with 10 µg/mL blasticidin, 250 µg/mL geneticin, 100 µg/mL zeocin and 100 µg/mL hygromycin for Jurkat cells and with 10 µg/mL blasticidin and 250 µg/mL geneticin for Raji cells.

The cellular assay was performed according to vendor instructions. Briefly, 2 days prior to the assay cells were seeded in medium without antibiotics at 5×10^5^ Jurkat cells/mL and at 4×10^5^ Raji cells/mL. On the day of the assay, the cells were pelleted and suspended in medium without antibiotics at 2.2×10^6^ Jurkat cells/mL and at 1.1×10^6^ Raji cells/mL. In 96-well round bottom Nunclon Delta plates (Thermo), per well, 90 µL of each cell suspension was added to 20 µL of antibody solution in PBS and the plates were incubated 6 hours at 37°C in a humidified incubator at 5% CO_2_. Each antibody concentration was assayed in triplicate on 3 independent plates. Each plate contained cells incubated with PBS without antibody as background control as well as cells treated with 50 nM dostarlimab used for signal normalization among different plates. After 6 hours incubation, 20 µL of the suspension of the co-cultured cells was transferred to 96-well white flat bottom TC plate (Greiner) and 50 µL of the working solution of the QUANTI1Luc 4 Lucia/Gaussia reagent (Invivogen) was added. Luminescence was measured immediately afterwards using a Tecan SPARK reader with 100 ms reading time. Luminescent signal measured in each well was expressed as % of signal elicited by 50 nM dostarlimab in the same plate and an average of results obtained in 3 independent plates was calculated.

### Thermal Unfolding

Thermal unfolding of SpyCatcher mutants was measured on a Prometheus nanoDSF device (Nanotemper) from 20°C to 90°C with a ramp rate of 2°C/minute.

## Acknowledgements

We thank our colleagues in the protein production and QC teams for generating and validating the reagents used in this study. We thank Achim Knappik for critically reading the manuscript.

## Conflict of Interest Statement

CH, MP, FY are authors of a patent application based on this work. All authors are employees of Bio-Rad AbD Serotec GmbH.

## Author Contributions

Conceptualization and design of experiments: CH, MP, FY. Performed experiments: HH, MP, SJK, MW, MC. Generated reagents: WP, SH. Writing: CH, MP, FY with input from all authors.

