## Supplementary information for "Engineered Reversible Inhibition of SpyCatcher Reactivity Enables Rapid Generation of Bispecific Antibodies"

[1]Bio-Rad AbD Serotec GmbH, Anna-Sigmund-Str. 5, 82061 Neuried, Germany

[*]These authors contributed equally

**Supplemental Figure 1: SpyCatcher3 cysteine scan – reactivity of cysteine mutants**

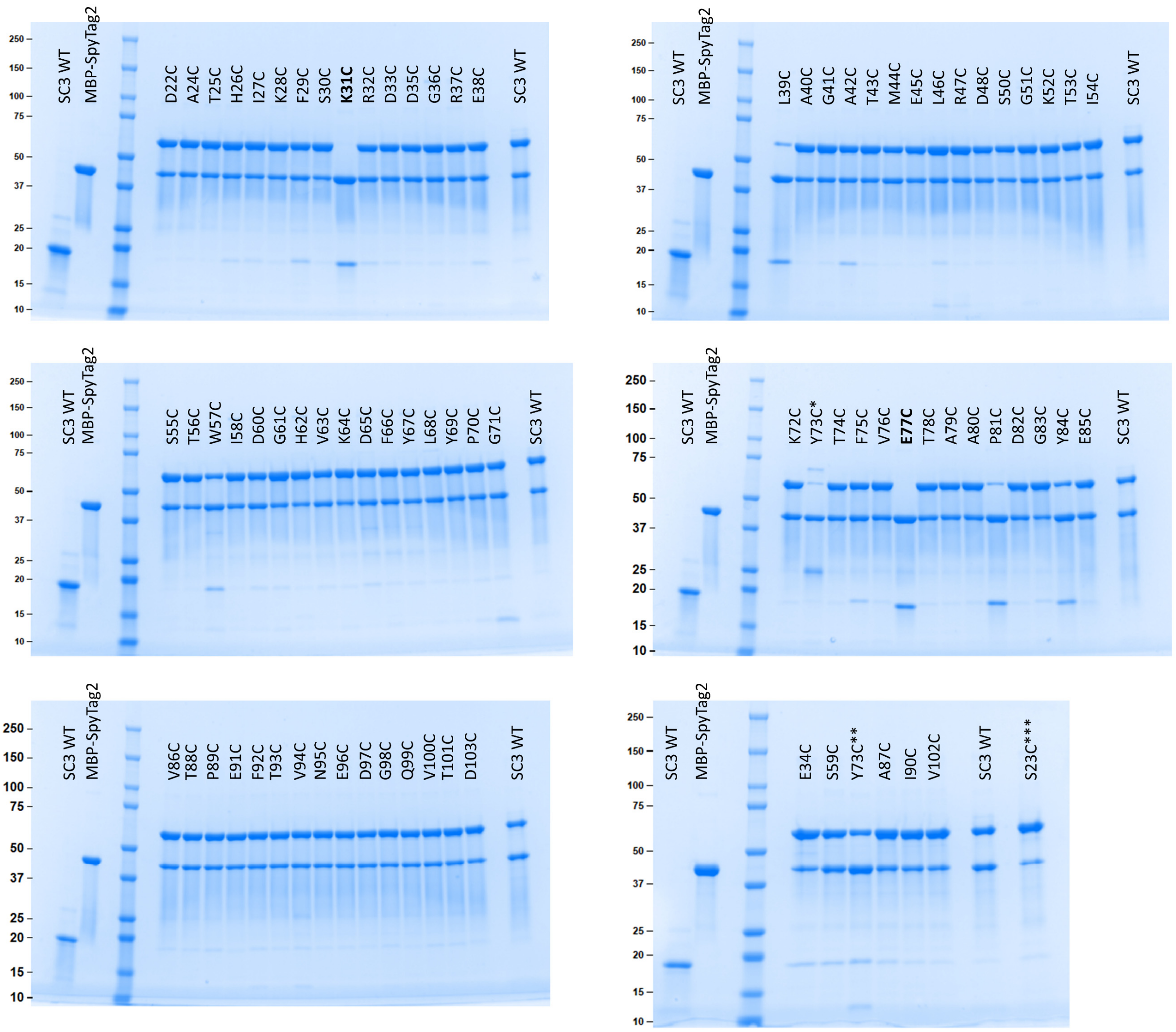


SDS-PAGE analysis of 81 SpyCatcher3 mutants and wildtype after coupling with MBP-SpyTag2. Input SpyCatcher3 wildtype and MBP-SpyTag2 are in two leftmost lanes of each gel. Reaction conditions: 4 µM SpyCatcher3 variant, 6 µM MBP-SpyTag2, 1 hour reaction time in PBS. The position of each mutant is indicated on the gel. The first purification of SpyCatcher3 Y72C (labeled with *) did not contain sufficient SpyCatcher and was therefore repeated (**). The S23C mutant (***) contains two additional cysteines, at the N-terminus and an S49C mutation. Reactive and catalytic residues are emphasized in bold print.

**Supplemental Figure 2: SpyCatcher3 cysteine scan – reactivity after modification with TNB**

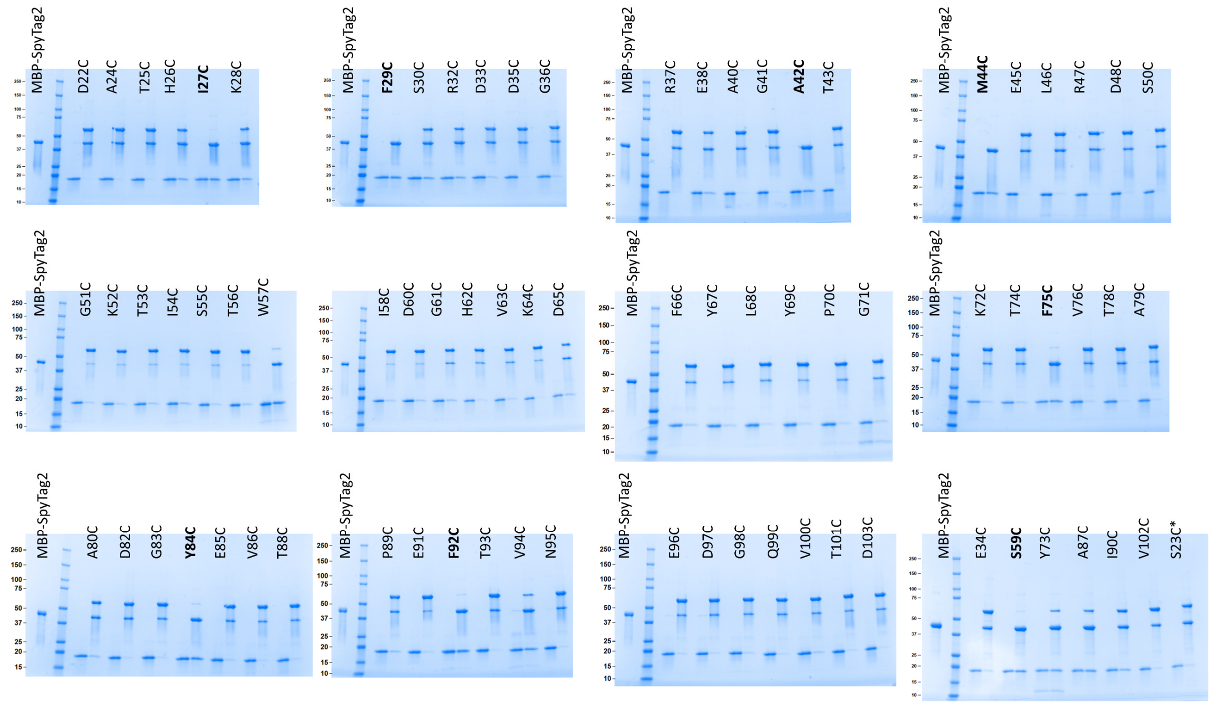


SDS-PAGE analysis of 79 different SpyCatcher3 cysteine mutants modified with Ellman's reagent. Input MBP-SpyTag2 is applied in the leftmost lane of each gel. Each SpyCatcher mutant is loaded in two sequential lanes, as input and as product of reaction with MBP-SpyTag2 (4 µM SpyCatcher3 mutant, 6 µM MBP-SpyTag2, 1.5 hours reaction time). Each pair of lanes is labeled with the mutant name. SpyCatcher3 mutants for which TNB modification led to inhibition of SpyTag coupling are highlighted in bold. The S23C mutant (*) contains two additional cysteines, at the N-terminus and an S49C mutation.

**Supplemental Figure 3: SpyCatcher3 cysteine scan – reversibility of inhibition by reduction**

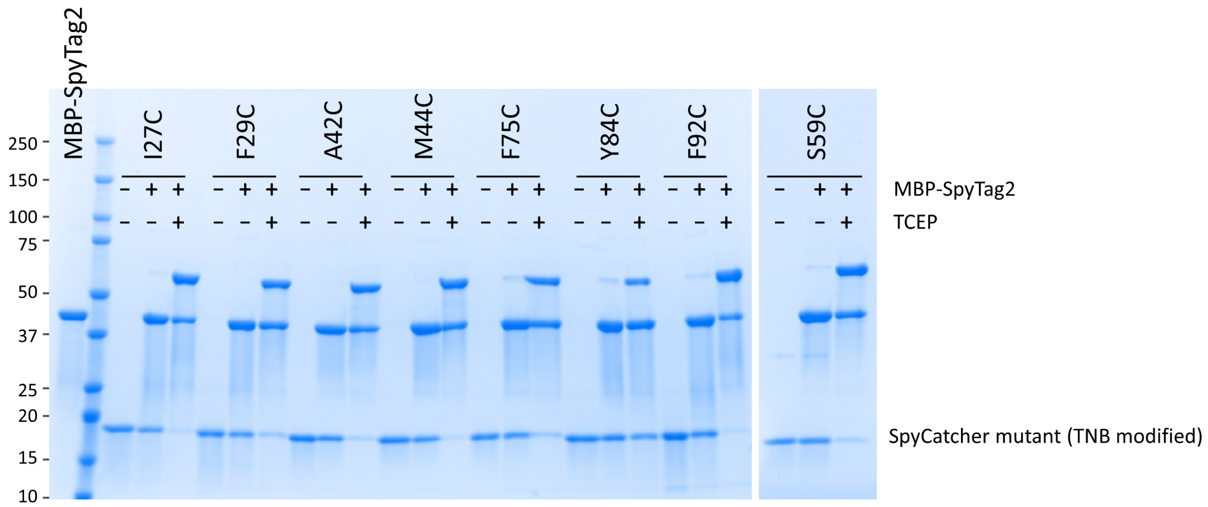


SDS-PAGE analysis of 8 SpyCatcher3 cysteine mutants for which SpyTag reactivity was inhibited by modification with TNB. Each mutant is loaded in three sequential lanes, as input in its TNB modified form, mixed with MBP-SpyTag2, and mixed with MBP-SpyTag2 in presence of 10 mM TCEP. The name of the respective SpyCatcher3 mutant is indicated above the corresponding gel lanes. Reaction conditions of SpyTag coupling: 4 µM SpyCatcher3 mutant, 6 µM MBP-SpyTag2, 1.5 hours reaction time in PBS.

**Supplemental Figure 4: Effectiveness of reversible inhibition with TNB and HPDP-biotin**

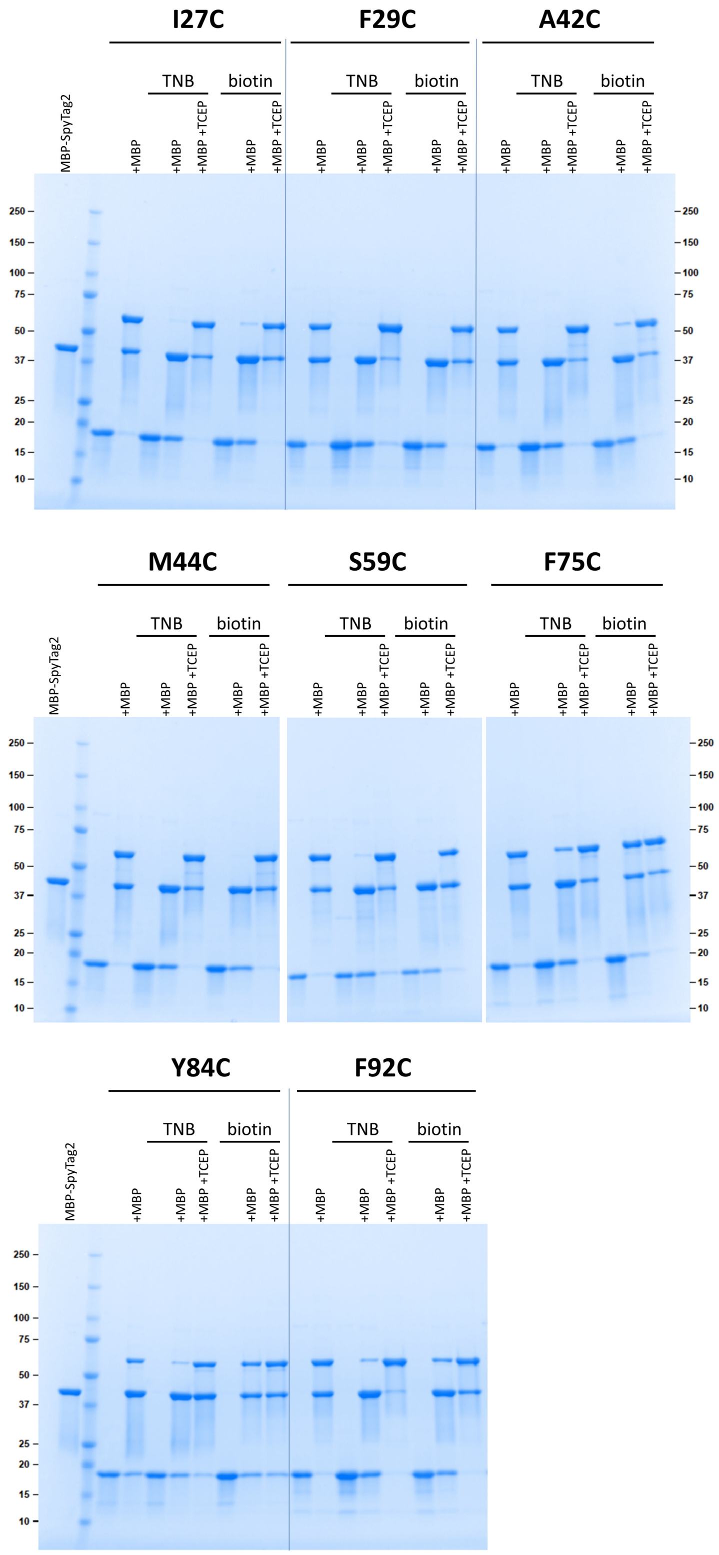


SDS-PAGE analysis of 8 SpyCatcher3 cysteine mutants without disulfide modification, modified with TNB and modified with HPDP-biotin, conjugated with MBP-SpyTag2 in the absence or presence of reducing agent. Each mutant is loaded in 8 sequential lanes, precise reaction conditions are indicated above each lane. Dependent on the reaction conditions, following concentrations of reactants were used: 4 µM SpyCatcher3 mutant, 6 µM MBP-SpyTag2 (also abbreviated as MBP in the gel labels), 5 mM TCEP. In all cases reaction time was 1.5 hours.

**Supplemental Figure 5: Unmodified SpyLock reaction time course**

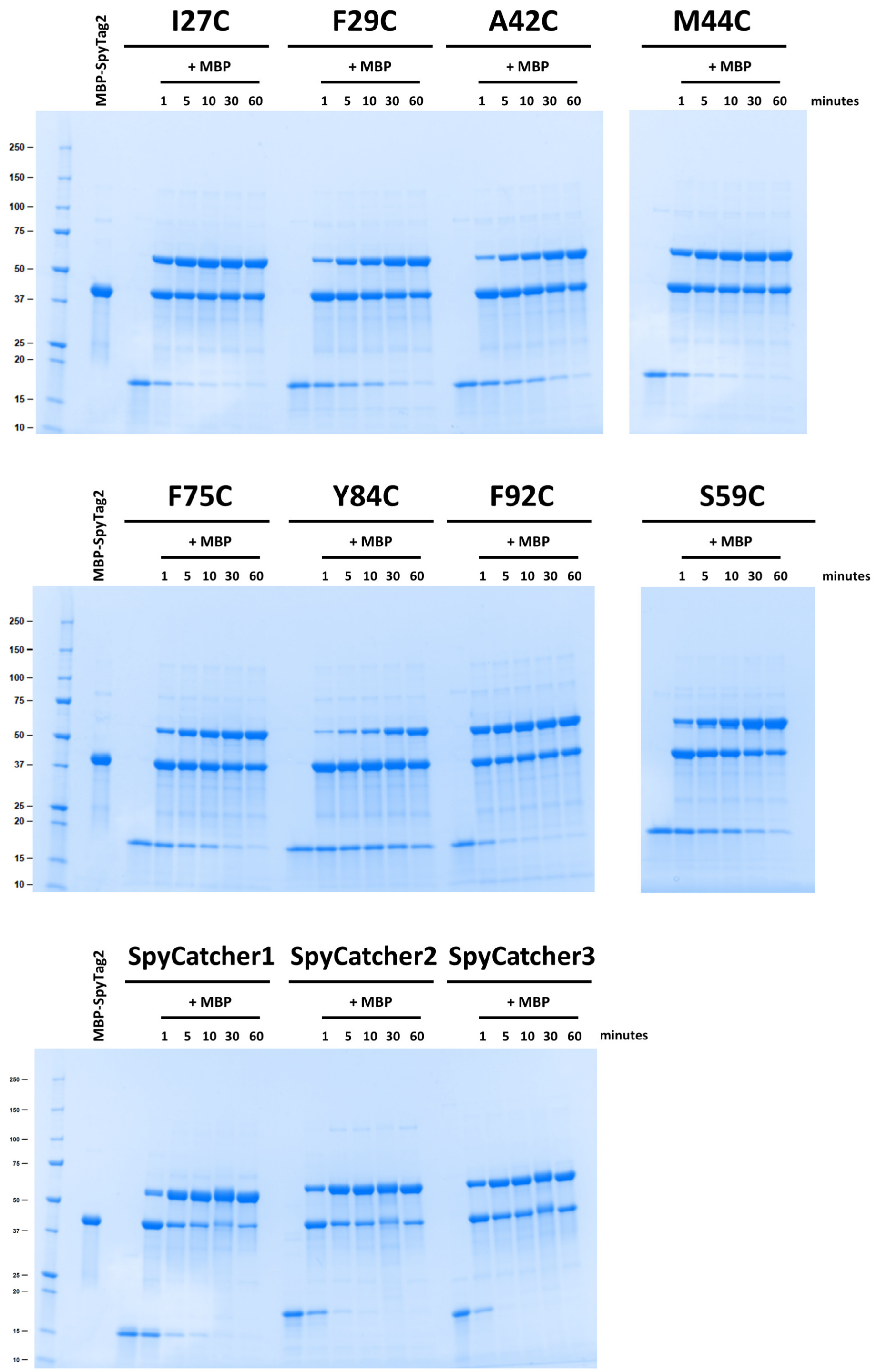


SDS-PAGE analysis of 8 SpyCatcher3 cysteine mutants and wildtype SpyCatcher1-3 reacting with MBP-SpyTag2. Reaction conditions: 4 µM SpyCatcher variant, 6 µM MBP-SpyTag2 (also abbreviated as MBP in gel labels). Reaction time is indicated above the corresponding lane.

**Supplemental Figure 6: Double band formation of BiCatcher3 after SpyTag coupling**

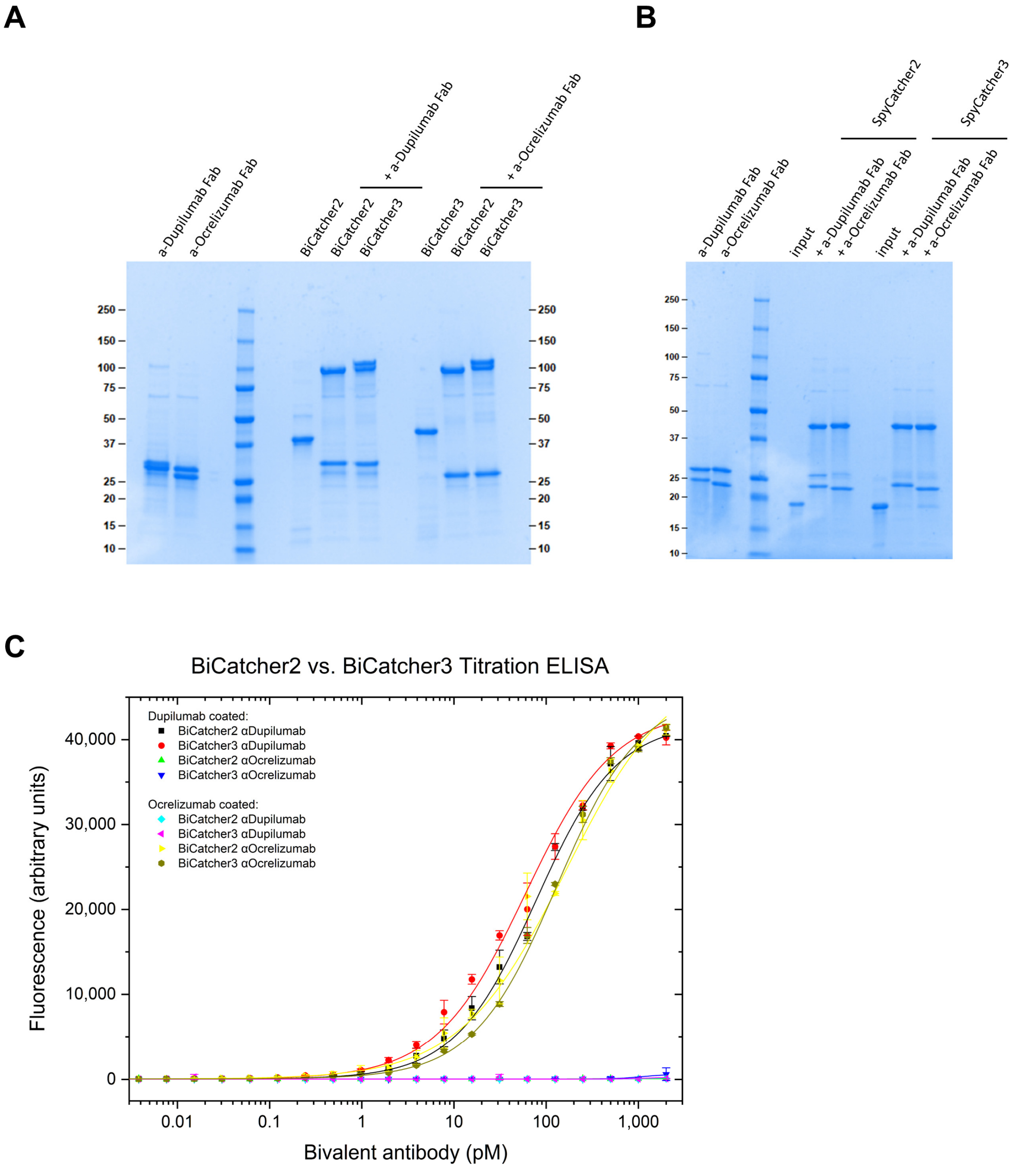


We observed an interesting aspect of producing bivalent constructs when using BiLock based on SpyCatcher3. The BiLock construct as well as BiCatcher3 coupling products migrated as double bands in SDS-PAGE when fully coupled, while fully coupled monomeric SpyCatcher3 does not form double bands. Bivalent constructs based on SpyCatcher1 and 2 do not exhibit this phenomenon. While this observation is at first glance reminiscent of the deamidation occurring in SpyCatcher1 and 2[1], a simple posttranslational modification cannot be the cause here, as the monovalent starting materials and monovalent coupling products do not exhibit double bands in SDS-PAGE. Given the limited number of amino acid differences between SpyCatcher2 and SpyCatcher3 — only five in total — one of these substitutions, possibly the additional proline, is likely responsible for the observed occurrence of double bands in dimeric constructs, which otherwise show a single band when reacting as individual monomeric proteins. In any case, there is no observable difference in affinity between bivalent constructs based on SpyCatcher2 and 3, therefore the difference is irrelevant for antibody screening.

A) Non-reducing SDS-PAGE gel of coupling reaction products of BiCatcher2 (SpyCatcher2-SpyCatcher2 fusion) and BiCatcher3 (SpyCatcher3-SpyCatcher3 fusion) with two different Fabs along input controls.

B) Reducing SDS-PAGE gel of coupling reaction products of monomeric SpyCatcher2 and monomeric SpyCatcher3 with two different Fabs along input controls.

C) Titration ELISA comparing the sensitivity and specificity of BiCatcher2 and BiCatcher3 bivalent constructs generated with anti-dupilumab and anti-ocrelizumab Fabs. Dupilumab or ocrelizumab were coated, as indicated. Stated bivalent constructs were titrated onto the plate and HRP-conjugated anti-His-Tag antibody was used for detection. Error bars represent standard deviation of 2 measurements. Logistic fit was applied.

**Supplemental Figure 7: Effect of TCEP incubation on antigen binding**

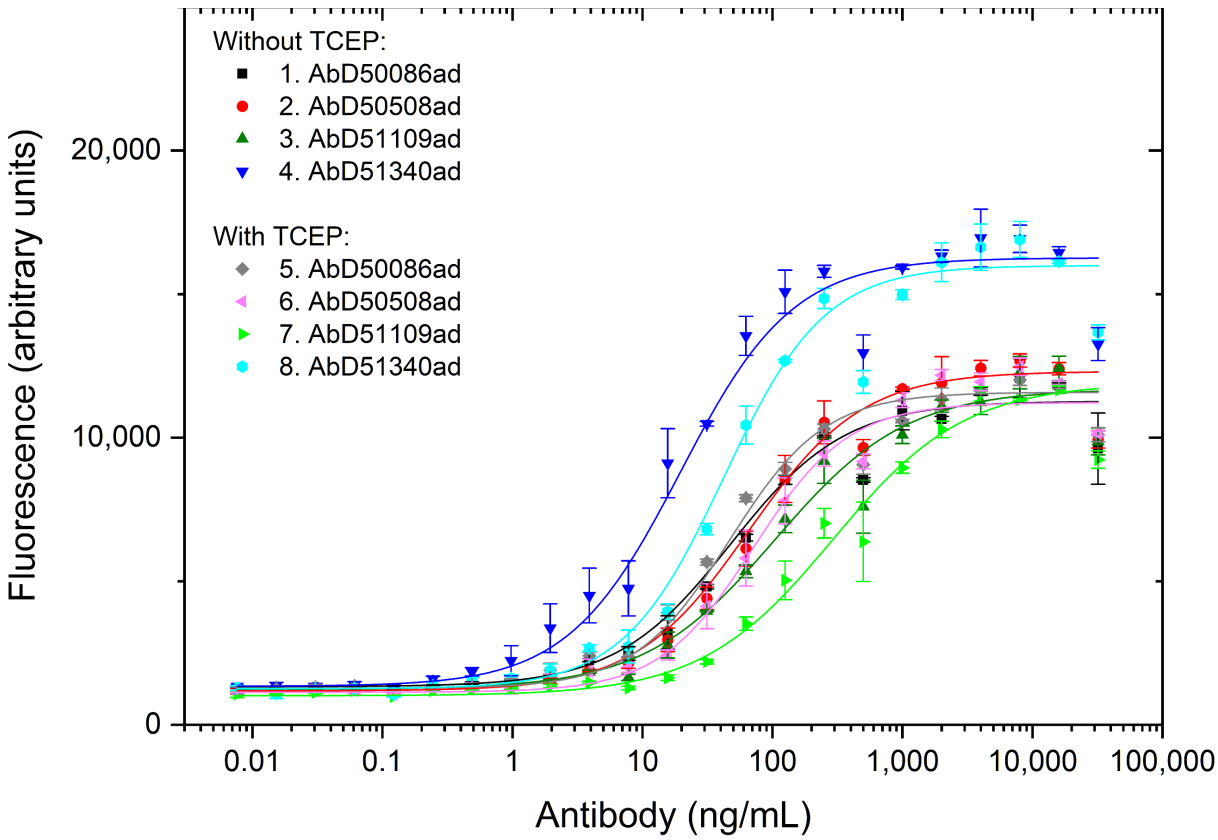


Titration ELISA of 4 different anti-GFP Fabs: AbD50086ad, AbD50508ad, AbD51109ad, AbD51340ad, after 1 week of incubation at 4°C with or without 5mM TCEP. mGFP was coated, anti-GFP Fabs were titrated and detection was performed with anti-Fab-HRP. Error bars represent standard deviation of 2 measurements. Logistic fit was applied.

### Supplemental Figure 8: Bispecific cell staining assay

A

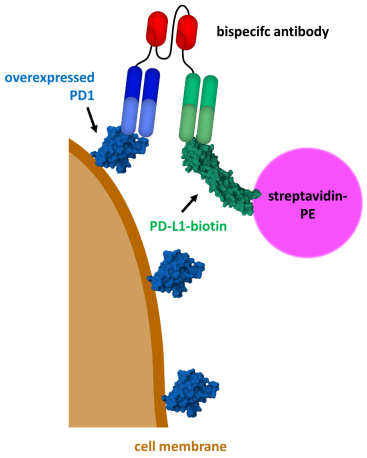


B

| Symbol | Antigen | International nonproprietary name |
|---|---|---|
| A | PD-L1 | atezolizumab |
| B | PD-L1 | avelumab |
| C | PD-L1 | durvalumab |
| 1 | PD-1 | cemiplimab |
| 2 | PD-1 | dostarlimab |
| 3 | PD-1 | nivolumab |
| 4 | PD-1 | pembrolizumab |

C

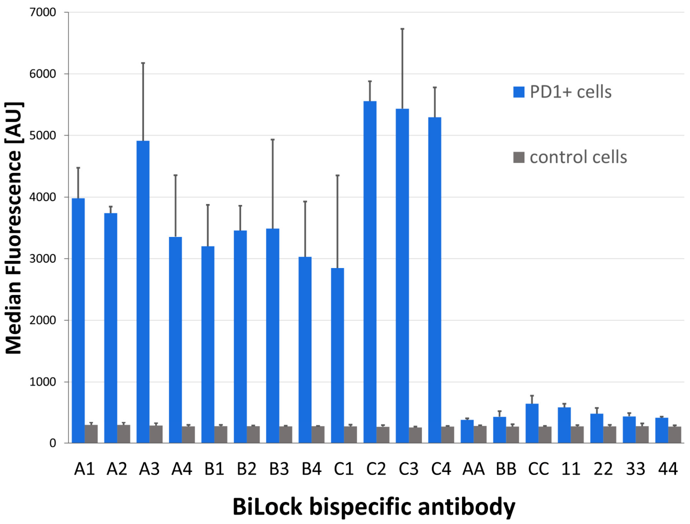


A) Schematic representation of the bispecific cell staining assay: PD-1-overexpressing HKB11 cells are incubated consecutively with BiLock-based bispecific antibodies or control antibodies, PD-L1-biotin and then stained with streptavidin-PE and analyzed by flow cytometry.

B) List of therapeutic anti-PD-L1 (antibodies A-C) and anti-PD-1 (antibodies 1-4) antibodies used in the cell staining assay. All antibodies were expressed as SpyTagged Fabs and used for construction of BiLock bispecific reagents. To exemplify the nomenclature of the obtained products: antibody 'B4' denotes bispecific avelumab-BiLock-pembrolizumab whereas '33' denotes monospecific nivolumab-BiLock-nivolumab.

C) Median fluorescence from cell staining of PD-1-overexpressing cells or control cells. Error bars are standard deviations of triplicate measurements.

**Supplemental Figure 9: Stability of BiLock (SpyCatcher3-SpyCatcher3 S59C)**

### BiLock-biotin:

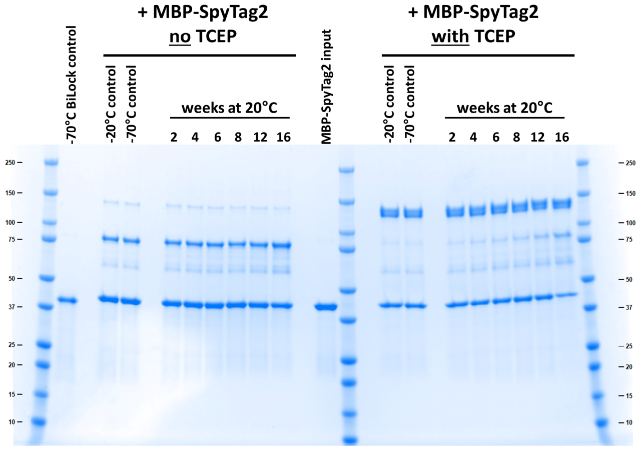


### BiLock-TNB:

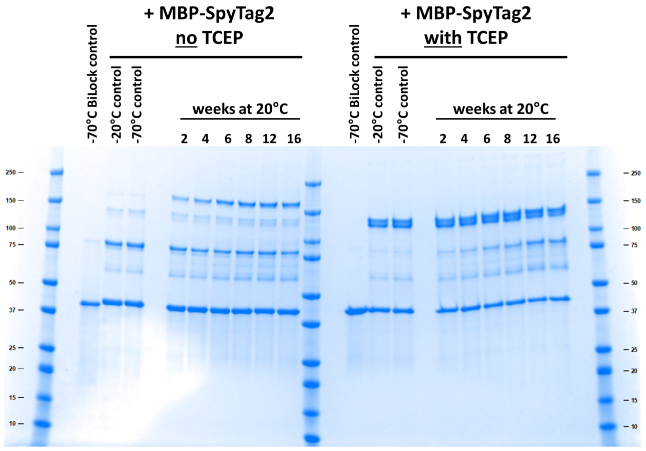


BiLock modified with HPDP-biotin or DTNB was incubated for 16 weeks at 20°C and samples were taken at regular intervals and stored at -70°C. Then, BiLock samples collected over 16 weeks were used for coupling with MBP-SpyTag2, which was performed either in absence or in presence of 20 mM TCEP. The coupled samples were applied on a non-reducing SDS-PAGE gel. BiLock samples stored for 16 weeks at -20°C and -70°C were used as controls. Reaction conditions: 5 µM BiLock, 12 µM MBP-SpyTag2, 1 hour at RT. Unconjugated BiLock and MBP-SpyTag2 input samples were also included. Left panel: samples of BiLock blocked with HPDP-biotin, right panel: BiLock samples blocked with DTNB.

**Supplemental Figure 10: SpyLock-suitable positions in SpyCatcher spacefilling model**

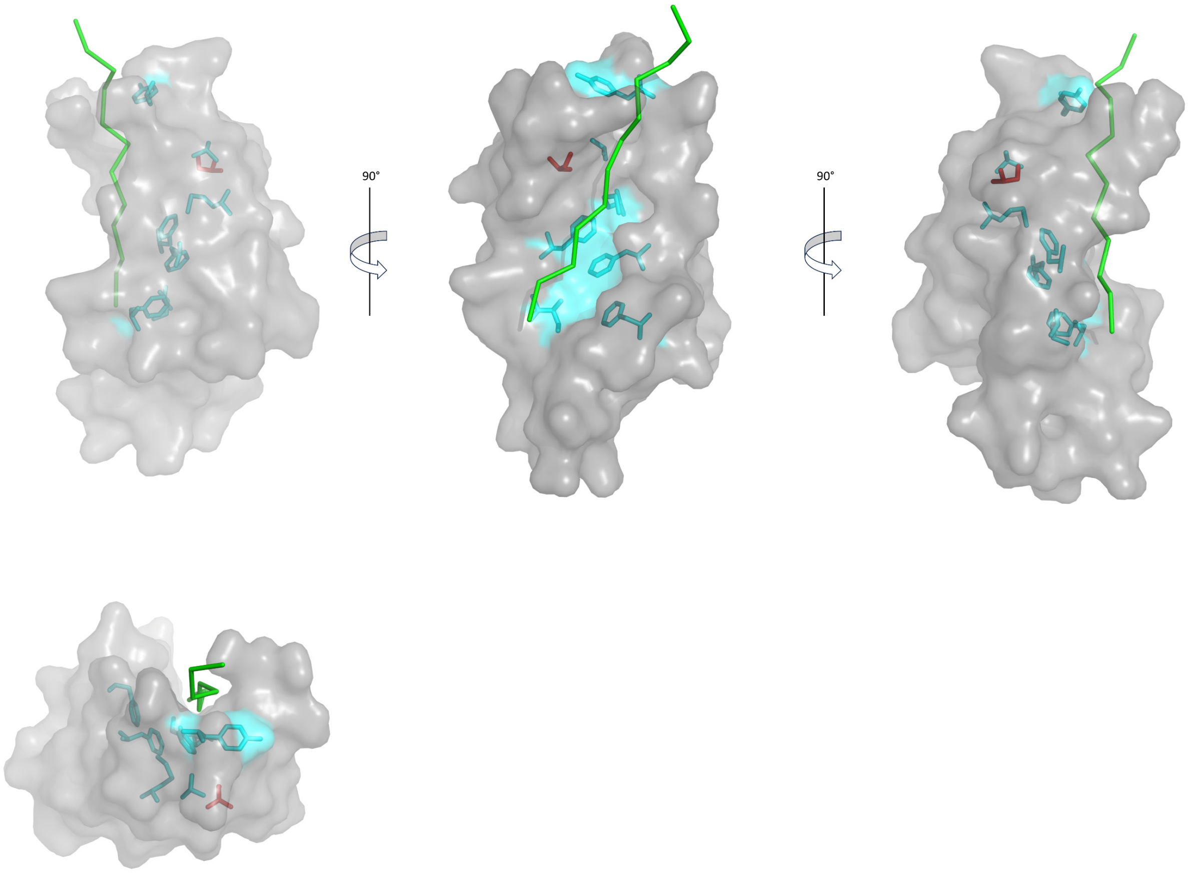


Different angles of showing the 8 identified SpyLock-suitable positions within SpyCatcher with respect to the SpyTag (green ribbon) binding groove. Structures generated from PDB 4MLI[2]. S59 is highlighted in red, the other positions in cyan.
